# Obesity induces phenotypic switching of gastric smooth muscle cells through the activation of the PPARD/PDK4/ANGPTL4 pathway

**DOI:** 10.1101/2024.10.27.620509

**Authors:** Sanaa Dekkar, Kamilia Mahloul, Amandine Falco, Karidia Konate, Romane Pisteur, Sarah Maurel, Laurent Maimoun, Norbert Chauvet, Prisca Boisguérin, David Nocca, Ariane Sultan, Florian Pallot, Guillaume Walther, Nicolas Cenac, Cyril Breuker, Sandrine Faure, Pascal de Santa Barbara

**Affiliations:** PHYMEDEXP, University of Montpellier, INSERM, CNRS, Montpellier, France; IRSD, University of Toulouse, INSERM, INRAe, ENVT, UPS, Toulouse, France; Department of Nuclear Medicine, University Hospital of Montpellier, Montpellier, France; CIC1411, INSERM, University Hospital of Montpellier, Montpellier, France; Department of Digestive Surgery, University Hospital of Montpellier, Montpellier, France; Department of Nutrition-Diabetes, University Hospital of Montpellier, Montpellier, France; LAPEC, University of Avignon, Avignon, France; Department of Clinical Pharmacy, University Hospital of Montpellier, Montpellier, France

## Abstract

**Objective:** Clinical research has identified stomach dysmotility as a common feature of obesity. However, the specific mechanisms driving gastric emptying dysfunction in patients with obesity remain largely unknown. In this study, we investigated potential mechanisms by focusing on the homeostasis of gastric smooth muscle, using tissue samples from both patient and mice, as well as human cell culture system.

**Design:** Tissue samples were collected from patients who underwent sleeve gastrectomy for obesity, as well as from control subjects with standard weight who were undergoing treatment for gastric or esophageal carcinoma. Stomach tissues were harvested from mice after 12 weeks on a High-Fat Diet (HFD). Differentiated human gastric smooth muscle cells (SMCs) were treated with lipids, siRNA-peptide-based nanoparticles and/or pharmaceutical compounds. All experimental conditions were analyzed for their effects on SMC differentiation using stage-specific smooth muscle markers. Additionally, lipidomic and RNA sequencing analyses were performed on human gastric SMC cultures. The findings were assessed in patients with obesity to evaluate their clinical relevance.

**Results:** The smooth muscle layers in gastric tissue from both patients with obesity and mice fed on a HFD exhibited altered differentiation status. Treatment of differentiated human gastric SMCs with lipids phenocopies these alterations and is associated with increased expression of *PDK4* and *ANGPTL4*. Inhibition of PDK4 or ANGPTL4 upregulation prevents these lipid-induced modifications. Mechanistically, lipid treatment activates PPARD, which regulates *PDK4* and *ANGPTL4* upregulation, leading to SMC dedifferentiation. Notably, *PDK4* and *ANGPTL4* levels correlate with immaturity and alteration of gastric smooth muscle in patients with obesity.

**Conclusion:** Obesity triggers a phenotypic change in gastric SMCs, driven by the activation of the PPARD/PDK4/ANGPTL4 pathway. These mechanistic insights offer potential biomarkers for diagnosing stomach dysmotility in patients with obesity.

## INTRODUCTION

Obesity is a prevalent chronic condition affecting approximately 13% of the adult population in the world, and constitutes a major and widespread global public health issue [1]. It results from mechanisms such as an imbalance between energy intake and expenditure, eating disorders and is strongly associated with various medical conditions, including type 2 diabetes, cardiovascular diseases. There is also strong evidence for an association of obesity with several cancer types (endometrial, postmenopausal breast, prostate, and renal) [2,3]. Individuals with obesity often experience gastrointestinal (GI) issues including gastroesophageal reflux, nonalcoholic fatty liver disease, and an elevated risk of colorectal and esophageal adenocarcinoma. Furthermore, functional GI disorders like dyspepsia, inflammatory bowel syndrome, constipation and diarrhea are frequently observed in these patients [4–7]. Despite their prevalence, a comprehensive understanding of GI complications associated with obesity remains insufficiently explored in both clinical practice and research.

Conflicting findings have been reported in studies investigating the impact of obesity on GI motility, likely due to the inherent regionalization and segmentation of the GI tract along the rostro-caudal axis, which complicates the assessment of obesity’s effects on GI function [8,9]. In line with the stomach’s role—alongside the brain—in regulating food intake, gastric emptying and satiety mechanisms, studies conducted on genetically and diet-induced obese mice have demonstrated significant alterations in gene expression in the stomach rather than in the small intestine or colon [9,10]. Several studies reported accelerated gastric emptying in patients with obesity, potentially leading to disrupted satiety signaling, to increased food consumption and contributing to the pathogenesis of obesity and its related complications [11–14]. Supporting this hypothesis, recent findings have associated faster gastric emptying with significant weight gain in younger adults [15].

Gastric emptying is a complex functional mechanism that requires the generation and conduction of regular depolarizing potential (slow waves) in the smooth muscle, events initiated by the interstitial cells of Cajal (ICC) which are electrically coupled to the smooth muscle cells (SMC) [16–19]. Interestingly, the stomach of early-induced obese mice shows increased proliferation and density of ICCs, contributing to the accelerated gastric emptying observed in these obese mice [20]. However, the impact of obesity on the gastric smooth muscle has been undervalued, even though disturbances in its activity can result in motility issues [21–25], possibly due to an incomplete understanding of GI smooth muscle physiology.

Digestive SMCs originate from LIX1-positive mesenchymal progenitors through a two-step process [26,27]. These progenitors first undergo a determination program characterized by the expression of MYOCARDIN, a cofactor of the transcription factor SRF (Serum Response Factor), which regulates the expression of smooth muscle actin proteins, including gamma (γSMA) and alpha (αSMA) smooth muscle actins [28]. Following this, determined SMCs enter a more differentiated state, marked by cell elongation and the expression of CALPONIN1 and SM22, two actin-binding proteins involved in smooth muscle contractility [23]. As SMCs differentiate, they acquire the capacity to generate force and contract, facilitated by the presence and organization of smooth muscle actin and myosin proteins. While skeletal muscle contains satellite cells that respond to growth and injury, a comparable reservoir of cells presenting pluripotency capacities has not been identified in smooth muscles. Instead, SMCs possess the capacity to dedifferentiate upon stimulation, revert to a synthetic mesenchymal phenotype, and re-enter the cell division cycle to proliferate [23,29]. This physiological process is crucial during the growth of the pediatric intestine smooth muscle [30]. However, an imbalance in plasticity favoring the mesenchymal progenitor state is thought to contribute to numerous diseases, including primary visceral myopathy, inflammatory bowel disease, diabetes, and metabolic disorders [24,31], ultimately leading to impaired GI motility [22,23].

In this study, we thoroughly examined the effects of obesity on gastric homeostasis, with a focus on muscular changes. Analysis of tissues from patients with obesity and diet-induced obesity mouse model revealed alterations in SMC differentiation and plasticity. Using a novel *in vitro* model of differentiated human gastric SMCs, we demonstrated that lipids act directly on SMC plasticity and identified the PPARD/PDK4/ANGPTL4 pathway as a key player in this process.

## METHODS

A more detailed description of the methods used in this study is provided in the Supplemental Methods section.

### Patients

The study involved collecting gastric wall samples from two patient groups. The first group included 15 patients with obesity (body mass index (BMI) ≥ 35 kg/m²) who underwent sleeve gastrectomy between September and December 2018 (see supplemental table 1). The second group comprised 4 lean control patients with a BMI below 30 kg/m^2^ who underwent surgeries for gastric or esophageal epithelial tumors in January 2019 (see supplemental table 2). Participants were included regardless of their weight or diabetic status, and their biological data were collected. Samples were collected in accordance with the ethical guidelines of Montpellier University Hospital (France), with informed consent obtained from each patient, permitting the use of their tissue samples for research purposes. Micro-dissected smooth muscle fibers from the corpus of the stomach were analyzed using Western blotting and quantitative RT-PCR.

### High-Fat Diet mice treatment

Male C57BL/6J mice (10-11 weeks old) from Janvier Laboratories (Saint-Berthevin Cedex, France) were individually housed under controlled conditions with a 12-hour light-dark cycle and had unrestricted access to food and water. The study adhered to the European Parliament Directive 2010/63/EU and was approved by the local ethics committee of Marseille (Approval Number: 2020050612125728). Male mice were randomly divided into two groups (n=7 per group): one group received a normal chow diet (Control) and the other a High-Fat Diet (HFD) containing 60% fat (230-HFD). After 12 weeks of the diet, mice were assessed for changes in metabolism, and biochemistry.

### Human gastric smooth muscle cell and treatment

Human gastric SMCs (Innoprot Innovative, Spain) were cultured on collagen I-coated dishes (Corning) in Dulbecco’s Modified Eagle’s Medium (DMEM) with 10% fetal bovine serum (Sigma) and 1% penicillin/streptomycin [27]. Differentiation was induced over 14 days, with pharmacological treatments commencing after differentiation. For lipid treatments, SMCs were exposed to a 2% lipid mixture (Sigma) supplemented with BSA-complexed long-chain fatty acids (30 µM palmitic acid, and 30 µM oleic acid), or a 60 µM BSA (fatty acid-free preparation of BSA) as control for 3 or 7 days. In the PPARD antagonist experiments, SMCs were serum-starved for 24 hours and treated with 5 µM GSK0660 or 0.1% DMSO for 24 hours, followed by exposure to the lipid treatment with additional GSK0660 every 24 hours for 72 hours. For PPARD agonist experiments, SMCs were treated with 1 µM GW501516 or 0.1% DMSO for 24 or 72 hours. To inhibit lipid-induced *PDK4* and *ANGPTL4* expression, SMCs were treated with WRAP5-siRNA nanoparticles targeting *PDK4*, *ANGPTL4*, or a control siRNA for 2 hours, then cultured in a lipid treatment for 3 days. Analysis included immunofluorescence, Western blotting, quantitative RT-PCR, and assessments of cell viability and toxicity. Gastric SMCs were routinely screened for mycoplasm contamination using the MycoAlert® Detection Kit (Lonza).

### Transcriptomic sequencing and lipidomic analysis

The gene expression profile of human gastric SMC cultures treated with lipids, along with their respective controls, was assessed using high-throughput RNA sequencing. Differentially expressed genes were identified using the DESeq2 statistical method (MGX platform, Biocampus, Montpellier, France). The lipid composition of the treated SMC cultures was analyzed using an Agilent 1290 UPLC system coupled to a G6460 triple quadripole spectrometer (Agilent Technologies) for the ceramides and the sphingomyelins, and by gas-liquid chromatography on a Clarus 600 Perkin Elmer system using a Famewax RESTEK fused silica capillary columns for total fatty acid (fatty acid methyl esters), as previously described [32,33].

### Statistical analysis

Data are presented as means ± standard error of the mean (SEM). Statistical analyses were performed using GraphPad Prism 9.0 software (GraphPad, San Diego, CA) with the two-tailed Mann-Whitney test. For multiple comparisons within groups, the Kruskal-Wallis test followed by Dunn’s multiple comparisons test was employed. Pearson’s correlation test was used for correlating gene and protein expression. Statistical significance was set at P < 0.05. Details of the statistical tests used are provided in the legend of each figure.

## RESULTS

### Obesity is associated with gastric musculature alterations in humans and mice

To investigate potential changes in gastric musculature associated with obesity, we first analyzed 15 samples from patients who underwent laparoscopic sleeve gastrectomy for obesity, as well as 4 samples from lean patients associated with gastric or esophageal carcinoma (controls) (supplemental tables 1 and 2). Histopathology and immunohistochemistry focused on smooth muscle (CALPONIN1) and enteric neuronal (TUJ1) markers. While some patients with obesity showed discrete changes in the organization of the smooth muscle layer, most exhibited significant muscular alterations associated with enteric neuronal hyperplasia (figure 1A,B). No alterations were found in the control subjects. Overall, more than 66% of patients with obesity showed muscle alterations (figure 1A,B; supplemental Table 1; supplemental figure 1A). Next, we evaluated the differentiation status of gastric smooth muscle cells (SMCs) using specific SMC marker antibodies. Immunohistochemical analysis revealed a reduced level of CALPONIN1 in the musculature of patients with obesity compared to controls (Supplemental Figure 1B). Western blot analysis of extracts from dissected gastric smooth muscle fibers confirms a significant decrease in the expression levels of differentiated markers, such as SM22 and CALPONIN1 in patients with obesity compared to controls. In contrast, the expression level of γSMA, a marker defining SMC determination and identity, remained unchanged indicating a specific deregulation of the differentiation status of the SMCs (figure 1C,D). We then analyzed the gastric musculature in a mouse model of early induced obesity. Male adult mice were subjected to a HFD for 12 weeks, a period known to induce various obesity-related metabolic changes [34]. This dietary regimen effectively induced obesity, as evidenced by the appearance of multiple metabolic phenotypes commonly associated with the condition (supplemental figure 2). Similar to our observations in human samples, Western blot analysis revealed a significant reduction in Sm22 and Calponin1 levels in gastric extracts prepared from obese mice compared to controls (figure 1E,F). γSMA levels remained constant (figure 1E,F). These findings suggest a link between obesity and alterations in the differentiation status of gastric smooth muscle.

**Figure 1.**
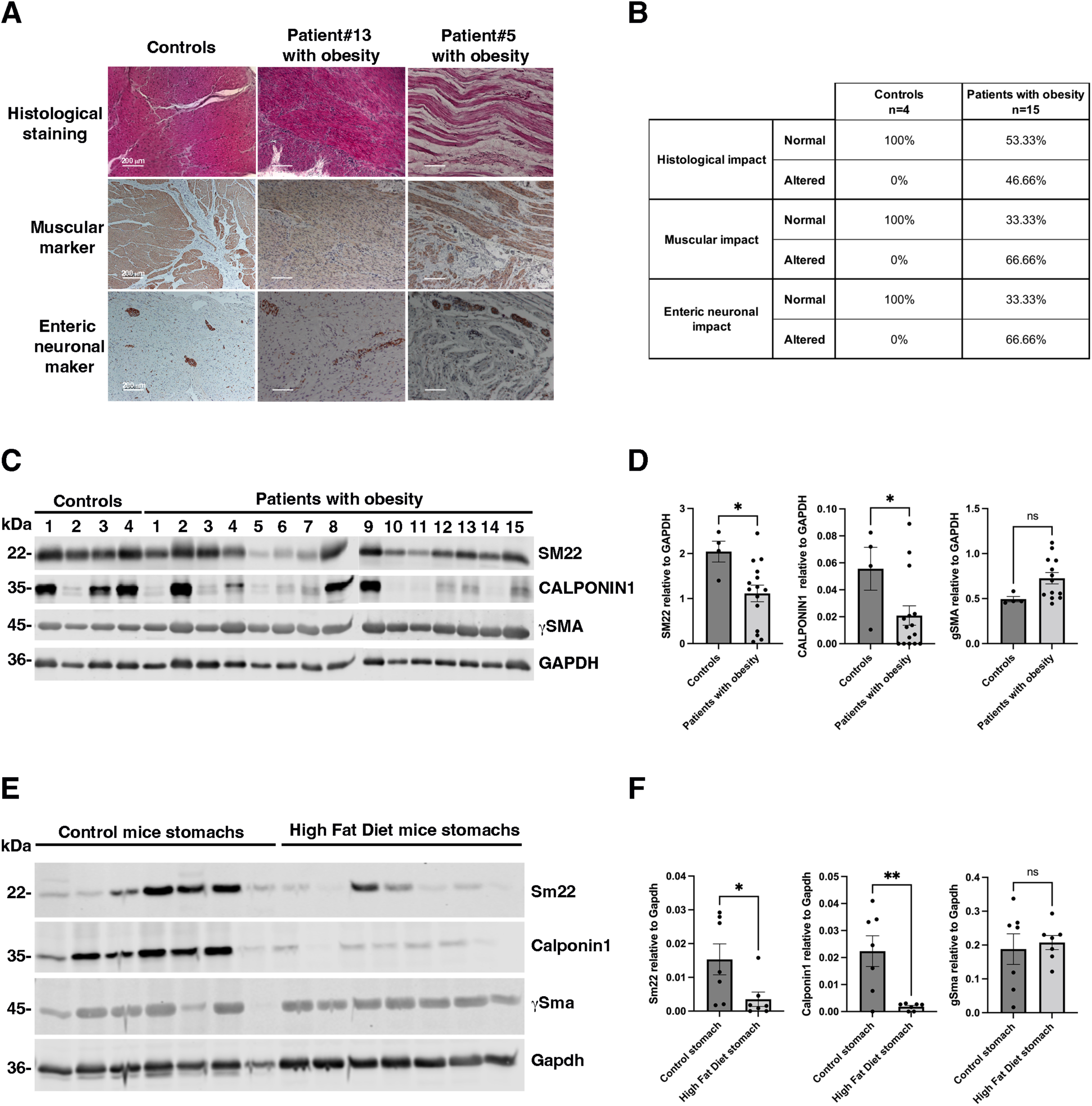
Evaluation of the differentiation status of human gastric smooth muscle in patients with obesity and in High fat diet (HFD) induced obese mice. (A) Representative Hematoxylin and Eosin staining (upper panels), immunohistochemistry staining of smooth muscle marker CALPONIN1 (medium panels) and of the enteric neuron marker TUJ1 (lower panels) of stomach sections from adult patients with obesity (n=15), and from lean adults with gastric epithelial cancer (controls; n=4). 10 of 15 patients with obesity exhibited alteration in the expression and organization of gastric smooth muscle and hyperplasia of myenteric enteric neurons. (B) Summary of the histological evaluation of stomach sections from adult patients with obesity (n=15) and adult controls (n=4) using Hematoxylin and Eosin staining, along with immunohistochemical staining for the smooth muscle marker CALPONIN1 and enteric neuron marker TUJ1. After staining, gastric smooth muscle was evaluated and scored independently for their organization into normal, and affected regions by three pathologists and consensus was reported. (C) Western-blotting of gastric smooth muscle fiber extracts from adult patients with obesity (n=15), and from controls (n=4) probed with antibodies directed against specific smooth muscle proteins (SM22, CALPONIN1, and γSMA) and against GAPDH as loading control. (D) Quantification of Western-blot assays (C) comparing extracts from adult patients with obesity (n=15) to extracts from controls (n=4). Data are presented as the mean ± SEM (two-tailed Mann–Whitney test; *P<0.05). Notably, the expression of SM22 and CALPONIN1 were found significantly lower in patients with obesity compared to controls (C, D). (E) Representative Western-blot image of whole stomach extracts from adult mice fed with a HFD for 12 weeks (n=7), and from controls (n=7) probed with antibodies directed against specific smooth muscle proteins (Sm22, Calponin1, and γSma) and against Gapdh as loading control. (F) Quantification of the Western-blot (E) comparing extracts from HFD stomach (n=7) to extracts from controls (n=7). Data are presented as the mean ± SEM (two-tailed Mann– Whitney test; *P<0.05; **P<0.01). The expression of Sm22 and Calponin1 was significantly lower in HFD compared to control condition (E, F).

### Lipid treatment altered gastric smooth muscle cell differentiation

Obesity has been shown to disrupt both intestinal epithelium homeostasis and the composition of the enteric nervous system [35,36]. These disruptions may contribute to the alteration observed in gastric smooth muscle, potentially through epithelial-mesenchymal or neuro-mesenchymal interactions [37,38]. To investigate whether components of the HFD can directly influence the gastric smooth muscle phenotype, we established an *in vitro* culture model of differentiated human gastric SMCs. *In vitro* studies on SMCs present challenges due to the spontaneous and uncontrolled phenotypic modulation of SMCs toward the synthetic and undifferentiated states [22–25,27,39]. Research on uterine SMCs has demonstrated that low-density culture conditions lead to reduced expression of contractile markers, while confluence decreases proliferative capacity and enhances the expression of differentiation marker expression [40]. In our study, culturing γSMA-positive human gastric SMCs to confluence over a period of 14 days induced the expression of SM22 and CALPONIN1, subsequently referred to as differentiated SMCs (Figure 2A; supplemental figure 3). Following differentiation, SMCs were exposed to lipid constituents of the HFD, including long-chain fatty acids supplemented with palmitate acid and oleate acid, as previously described [35]. Lipid treatment resulted in the rapid internalization of the lipids, as evidenced by Bodipy detection through confocal microscopy after 6 hours (Figure 2B). Prolonged intracellular localization of lipids within lipid droplets was further observed through transmission electron microscopy 7 days post-treatment (Figure 2C). Importantly, SMC viability remained unaffected by lipid treatment after both 3 and 7 days of exposure (Figure 2D). Liquid chromatography-tandem mass spectrometry lipidomic analysis of the SMC membrane revealed an increase in fatty acids, primarily polyunsaturated fatty acids, at both 3 and 7 days of treatment (Figure 2E, left panel; supplemental table 3). This increase was accompanied by a specific decrease in total ceramides with no changes observed in total sphingomyelins, suggesting a pre-metabolic condition (Figure 2E, middle and right panels; supplemental table 4). Next, we assessed the impact of lipid treatment on SMC homeostasis. Notably, lipid exposure was associated with a reduction in the protein levels of SM22 and CALPONIN1, while γSMA expression remained unchanged (Figure 2F,G). These findings highlight the direct role of lipids in promoting SMC dedifferentiation and enhancing cellular plasticity.

**Figure 2.**
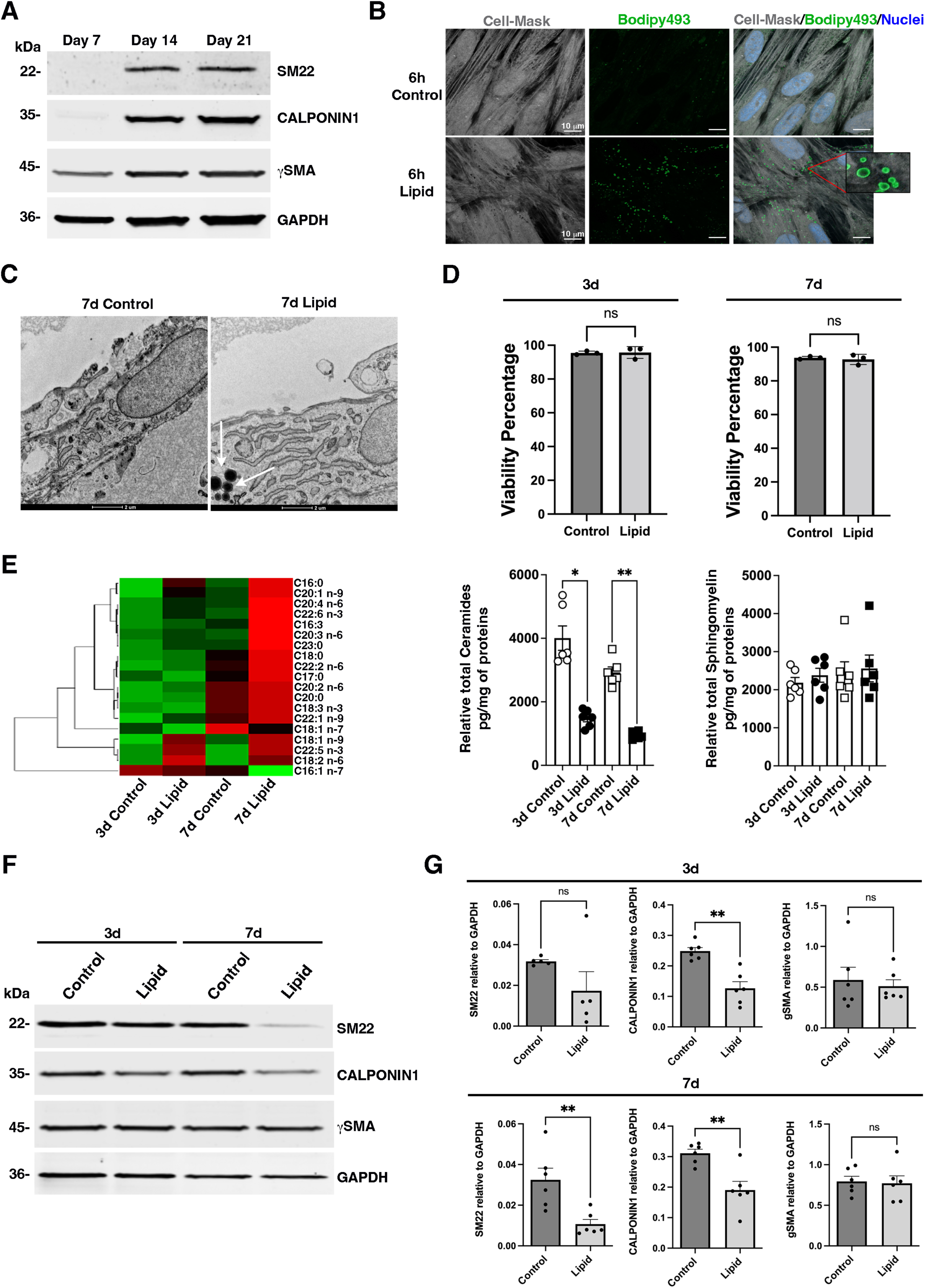
Impact of lipid treatment on human gastric smooth muscle cells. Human gastric smooth muscle cells (SMCs) were cultured on Collagen Type I plates over time to promote their differentiation (A, B). (A) Representative Western blot of human gastric SMC extracts from 7-, 14- and 21-days of culture probed with antibodies directed against specific smooth muscle proteins (SM22, CALPONIN1, and γSMA) and against GAPDH as loading control. (B) 14-day human gastric SMCs were incubated with lipid treatment for 6 hours, followed by staining with Bodipy493/503 and Cell-Mask Red stain 588/612 for 30 minutes, and nuclei staining with Hoechst. Scale bars: 10 µm. Numerous lipid structures within lipid-treated SMCs were detected with Bodipy staining (see zoomed insert) by confocal microscopy. (C) 14-day human gastric SMC cultures with or without lipid treatment for 7 days were analyzed by transmission electron microscopy. White arrows indicate the presence of lipid droplets observed only in the lipid-treated conditions Scale bars: 2 µm. (D) Cell viability of human gastric SMC cultures after 14 days with or without lipid treatment for an additional 3 or 7 days was evaluated by flow cytometry following differential DNA-binding dye staining. No significant impact on cell viability was observed after 3 days or 7 days of lipid treatment. Data are presented as the mean ± SEM and Mann-Whitney was applied (ns > 0.05). (E) Heat map of fatty acids (left panel) and levels of total ceramides (middle panel) and total sphingomyelins (right panel) quantified in SMCs with or without lipid treatment for an additional 3 and 7 days. For fatty acids, the data is shown in a matrix format: each row represents a single lipid, and each column represents a SMC treatment: control day 3, lipid day 3, control day 7, and lipid day 7. Each color patch represents the normalized quantity of lipid (row) in treated SMC (column), with a gradient from bright green (lowest) to bright red (highest). The pattern and length of the branches in the left dendrogram reflect the relatedness of the lipids. All data were normalized to the quantity of protein (pg/mg of proteins). Values are the mean ± SEM of n=6 samples. *P < 0.05 and **P < 0.01 (non-parametric Kruskall-Wallis test with Dunn’s multiple comparison test). (F) Representative Western blot of human gastric SMC extracts after 14 days of culture with or without lipid treatment for an additional 3 or 7 days probed with antibodies directed against specific smooth muscle proteins (SM22, CALPONIN1, and γSMA) and against GAPDH as loading control. (G) Quantification of Western blot assays (G) comparing extracts from lipid-treated SMCs to extracts from control SMCs. Data are presented as the mean ± SEM and two-tailed Mann–Whitney test was applied (*P < 0.05; ns > 0.05). CALPONIN1 expression was significantly reduced after 3 days, while SM22 was significantly lower only after 7 days of treatment (F, G).

### PDK4 and ANGPTL4 are essential in lipid-induced dedifferentiation of human gastric SMCs

To gain a deeper understanding of how lipid treatment mediates these effects, we conducted RNA sequencing analysis on human gastric SMCs after 3 and 7 days of lipid treatment. Using a significance cutoff of p < 0.01, we identified 139 up-regulated and 88 down-regulated genes compared to unstimulated SMCs at both time points (figure 3A; supplemental figures 4,5). Notably, *PDK4* and *ANGPTL4* were found to be strongly upregulated (figure 3B; supplemental figures 4,5). The stimulation of *PDK4* and *ANGPTL4* transcript level at 3 and 7 days of lipid treatment was confirmed by RT-qPCR (figure 3C,D). Importantly, when the concentration of lipid treatment is reduced by a factor of four, the upregulation of *PDK4* and *ANGPTL4* expression was no longer observed (supplemental figure 6A). Pyruvate dehydrogenase kinase family proteins (PDKs 1-4) inhibit glycolysis-dependent oxidative phosphorylation (OXPHOS) by inactivating pyruvate dehydrogenase complex, while angiopoietin-like proteins (ANGPTLs 1-8) are involved in various physiological and pathological functions related to tissue repair and homeostasis [41,42]. Among the PDK and ANGPTL family members, only *PDK4* and *ANGPTL4* were upregulated in response to lipid treatment (supplemental figure 6B,C).

**Figure 3.**
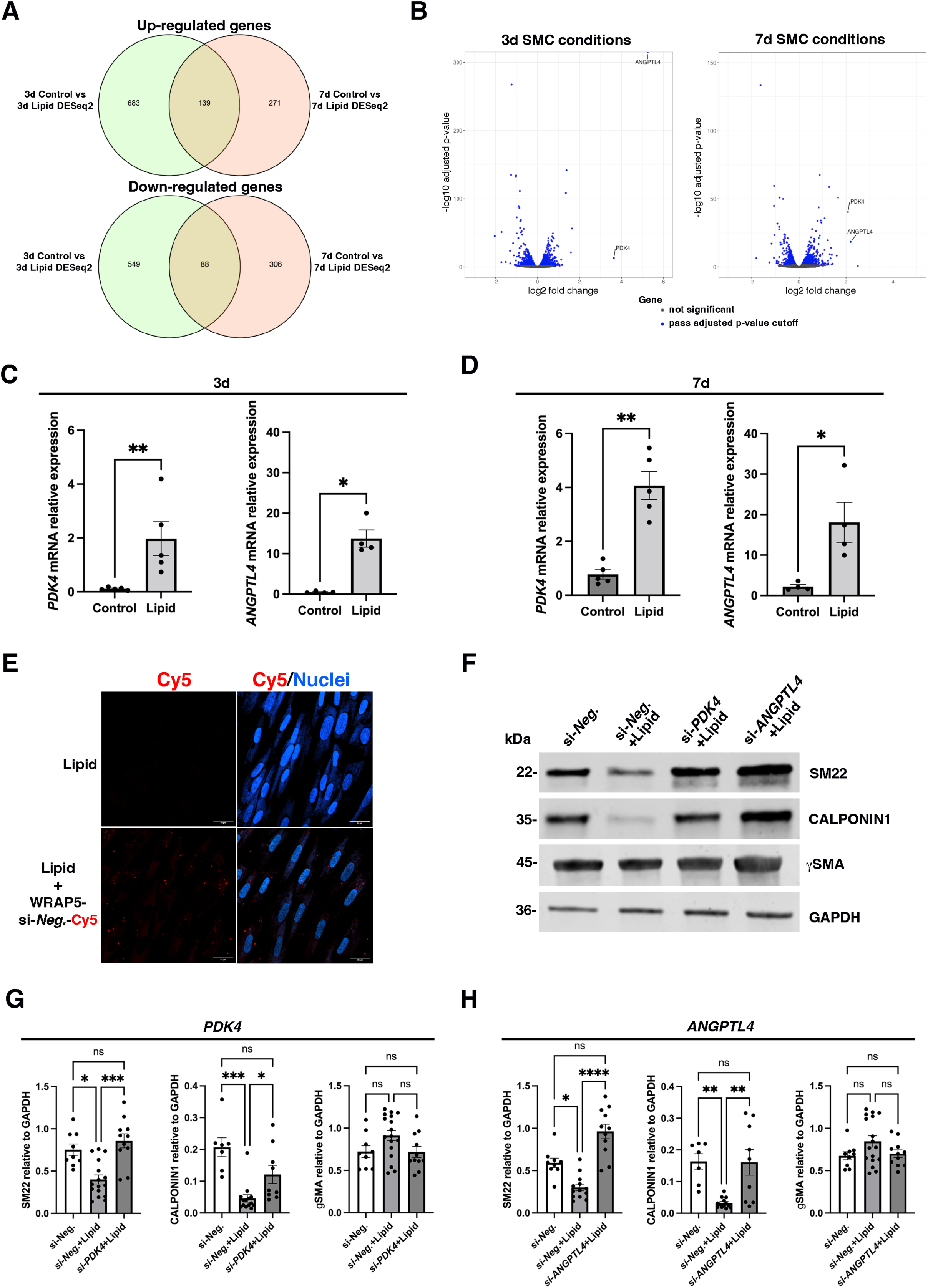
Identification and function of the molecular mechanism induced by lipid treatment in human gastric SMC. (A) Venn diagram depicting the number of genes up-regulated (upper panel) and down-regulated (lower panel) identified at 3 days and 7 days post-lipid treatment using a significance cutoff of p < 0.01. For both times, 139 genes were up-regulated, and 88 genes were down-regulated in response to lipid treatment. (B) Volcano plot showing differentially expressed genes between human gastric SMCs treated with lipid for 3 days (left panel) or 7 days (right panel) compared to untreated SMCs. Blue dots indicate genes with a significance cutoff of p < 0.01. Among them, upregulation of *PDK4* and *ANGPTL4* levels are found in both time conditions. (C) RT-qPCR of *PDK4* and *ANGPTL4* relative mRNA level in human gastric SMC cultures with or without lipid treatment for 3 days. Data were normalized to the house-keeping *HMBS* expression. Values are presented as the mean ± SEM of n=4-5 samples. Statistically significant differences are indicated as *P < 0.05 and ***P < 0.001 (two-tailed Mann–Whitney test). After 3 days of lipid treatment, *PDK4* and *ANGPTL4* levels were stimulated 13-fold and 29-fold, respectively. (D) RT-qPCR of *PDK4* and *ANGPTL4* relative mRNA level in human gastric SMC cultures with or without lipid treatment for 7 days. Data were normalized to the house-keeping *HMBS* expression. Values are the mean ± SEM of n=4-5 samples. Statistically significant differences are indicated as *P < 0.05 and ***P < 0.001 (two-tailed Mann–Whitney test). After 7 days of lipid treatment, *PDK4* and *ANGPTL4* levels were stimulated 5-fold and 8-fold, respectively. (E) 14-day human gastric SMC cultures were first incubated with WRAP-complexed Scrambled siRNA labeled with Cy5 (WRAP5+si-Neg.-Cy5) during 2 hours then cultures were treated with lipids. Following Hoechst staining, non-fixed cultures were directly analyzed using confocal microscopy. Scale bars: 24 µm. Many red dots detected with Cy5 staining were found in lipid-treated SMCs. (F) Representative Western blot of human gastric SMC extracts treated for 3 days with si-*Neg.* alone (control), si-*Neg.*+lipids, si-*PDK4*+lipids and si-*ANGPTL4*+lipids than probed with antibodies directed against specific smooth muscle proteins (SM22, CALPONIN1, and γSMA) and against GAPDH as loading control. (G) Quantification of Western blot assays comparing extracts treated for 3 days with si-*Neg.* alone (control), si-*Neg.*+lipids, and si-*PDK4*+lipids. Data are presented as the mean ± SEM (nonparametric Kruskall-Wallis test with Dunn’s multiple comparison test: *P < 0.05, ***P < 0.001, ns > 0.05). The expression of SM22 and CALPONIN1 decreased in lipid-treatet SMCs compared to the control condition; however, this decrease was abolished in the presence of si-*PDK4*. (H) Quantification of Western blot assays comparing extracts treated for 3 days with si-*Neg.* alone (control), si-*Neg.*+lipids, and si-*ANGPTL4*+lipids. Data are presented as the mean ± SEM (nonparametric Kruskall-Wallis test with Dunn’s multiple comparison test: *P < 0.05, **P < 0.01, ***P < 0.001, ns > 0.05). The expression of SM22 and CALPONIN1 decreased in lipid-treated SMCs compared to the control condition; however, this decrease was abolished in the presence of si-*ANGPTL4*.

To investigate the role of *PDK4* and/or *ANGPTL4* upregulation in the dedifferentiation of human gastric SMCs, we employed a method for cellular delivery of siRNA targeting these genes, avoiding lipid-based siRNA transfection systems. For this purpose, we utilized a 16-mer tryptophan- and arginine-rich amphipathic peptide (WRAP) [43], which forms nanoparticles with siRNA molecules, averaging 100 nm in size and exhibiting no cytotoxicity on human gastric SMCs (supplemental figure 7A-C). We found that Cy5-labeled WRAP5-siRNA nanoparticles were efficiently internalized by nearly all human gastric SMCs within 2 hours (figure 3E). Targeting *PDK4* or *ANGPTL4* with siRNAs effectively reduced the upregulation induced by lipid treatment (supplemental figure 7D), restoring SM22, CALPONIN1, and γSMA expression levels similar to those of unstimulated SMCs (figure 3F-H). Overall, our findings underscore the crucial role of PDK4 and ANGPTL4 as key regulators in driving the dedifferentiation of human gastric SMCs in the presence of lipids.

### PPARD regulates the induction of *PDK4* and *ANGPTL4* in lipid-treated gastric SMCs

We investigated the mechanisms by which lipid treatment induces the expression of PDK4 and ANGPTL4. Peroxisome proliferator-activated receptors (PPARα, PPARγ, and PPARβ/δ (also known as PPARD)) are ligand-activated transcription factors that regulate gene expression by forming heterodimers with RXRs and binding to PPAR response elements (PPREs) in the promoter regions of target genes [44,45]. These receptors can be activated by various ligands, including fatty acids [46]. Notably, PPARD directly binds to the PDK4 promoter, leading to its upregulation [47]. Bioinformatic analysis revealed conserved PPAR-RXRα binding sites in the promoters of *PDK4* and *ANGPTL4*, suggesting their regulatory role in response to lipid stimulation (figure 4A). Among the PPAR family, PPARD is the most abundantly expressed member of the PPAR family in human gastric SMCs (supplemental figure 8A). *PPARD* mRNA, along with that *PDK4* and *ANGPTL4* mRNA, is rapidly induced by lipids (supplemental figure 8B). PPARD nuclear localization is induced by lipid treatment (figure 4B). Furthermore, sustained activation of PPARD with its selective agonist GW501516 significantly enhanced *PDK4* and *ANGPTL4* mRNA expression, and to a greater extent than lipid treatment (figure 4C; supplemental figure 9). Treatment of SMCs with GW501516 resulted in a reduction of SM22 and CALPONIN1 levels after one day of treatment, while γSMA expression decreased after three days (figure 4D,E). Additionally, GSK0660, a potent and selective PPARD antagonist, inhibits the lipid-induced nuclear accumulation of PPARD (figure 4F). When combined to lipid treatment, GSK0660 blocked the upregulation of *PDK4* and *ANGPTL4* mRNA (figure 4G). These findings highlight the significant role of PPARD activity in regulating *PDK4* and *ANGPTL4* mRNA, which in turn drive the dedifferentiation of human gastric SMCs in response to lipid stimuli.

**Figure 4.**
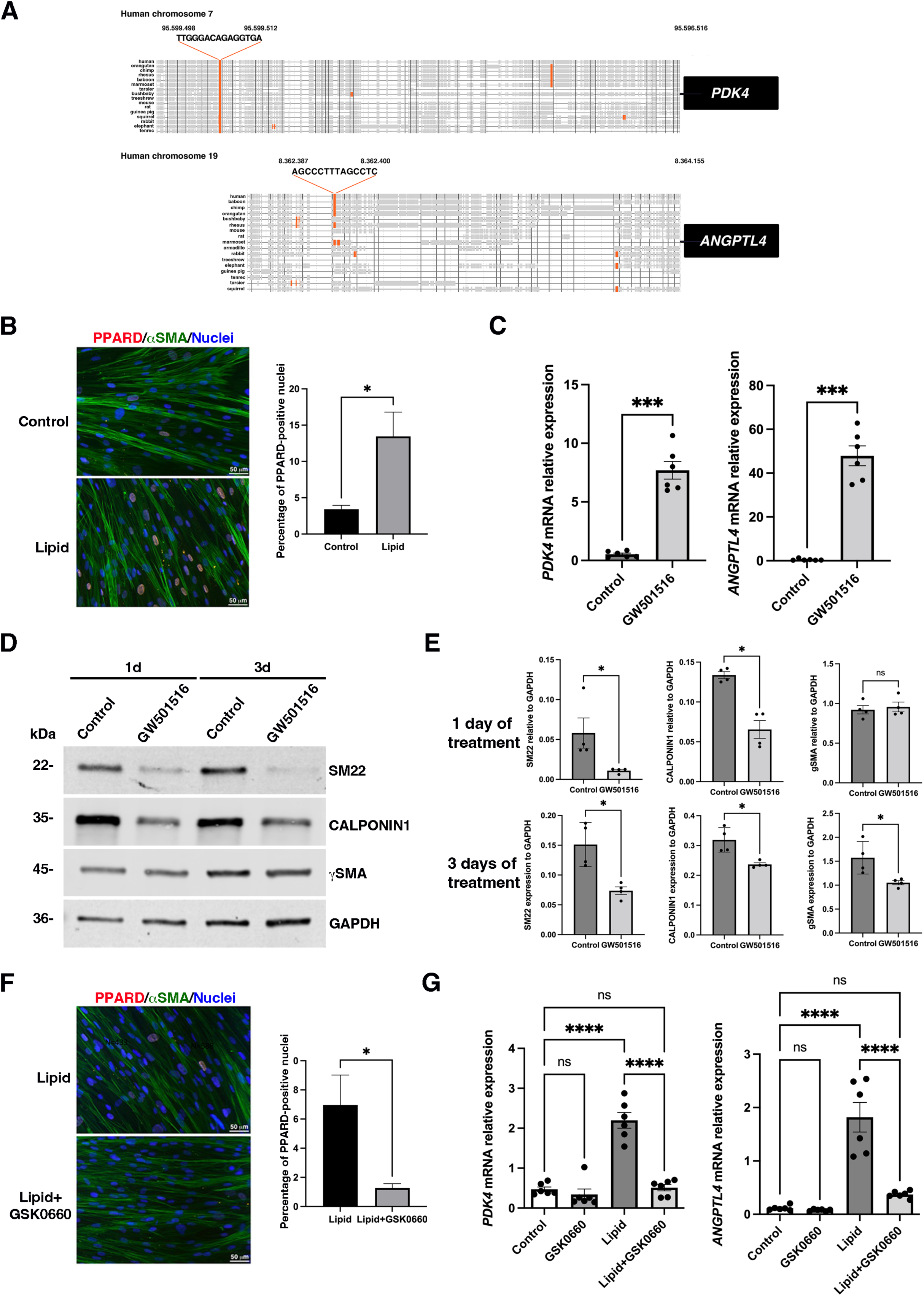
Regulation of the expression of *PDK4* and *ANGTPL4* mRNA expression under lipid treatment. (A) Promoter analyses of vertebrate *PDK4* and *ANGPTL4* genes using Contrav3 identified highly conserved PPAR-RXR binding sites across vertebrate species for *PDK4* and exclusively in primates for *ANGPTL4*. (B) Immunofluorescence analysis of human gastric SMC cultures treated with lipid for 2 days compared to untreated controls, stained for PPARD and αSMA. Nuclei were visualized using Hoechst staining. Scale bars: 50 µM. The right panel panel shows a 4-fold increase in the percentage of nuclear PPARD-positive SMCs following lipid treatment. (C) RT-qPCR of *PDK4* (left panel) and *ANGPTL4* (right panel) relative mRNA levels in human gastric SMC cultures treated for 2 days with GW501516 (PPARD agonist, 1 µM), compared to untreated SMC (Control). Data were normalized to the house-keeping *HMBS* expression. Values are the mean ± SEM of n=6 samples. ***P < 0.001 (two-tailed Mann– Whitney test). *PDK4* and *ANGPTL4* levels were stimulated 14.5-fold and 100-fold, respectively. (D) Representative Western blot of human gastric SMC extracts from 14 days of culture with or without GW501516 treatment for additive 1 day or 3 days probed with antibodies directed against specific smooth muscle proteins (SM22, CALPONIN1, and γSMA) and against GAPDH as loading control. (E) Quantification of Western blot assays (n=6) comparing extracts from GW501516-treated SMCs to extracts from control SMCs. Data are presented as the mean ± SEM and two-tailed Mann–Whitney test was applied (**P < 0.01, *P < 0.05, ns > 0.05). Significant reductions in SM22 and CALPONIN1 protein levels were observed after 1 and 3 days of treatment, with γSMA significantly lower only after 3 days (D, E). (F) Immunofluorescence analysis of human gastric SMC cultures treated for 1 day with lipids alone or in combination with GSK0660 (5 µM), stained for PPARD and αSMA. Nuclei were visualized using Hoechst staining. Scale bars: 50 µM. The right panel shows that lipid+GSK0660 treatment reduced the percentage of nuclear PPARD-positive SMCs by 6-fold. (G) RT-qPCR of *PDK4* (left panel) and *ANGPTL4* (right panel) relative mRNA levels in human gastric SMC cultures treated for 3 days with GSK0660 (PPARD antagonist, 5 µM), with lipid, and with lipid+GSK0660 compared to untreated SMC (Control). Data were normalized to the house-keeping *HMBS* expression. Values are the mean ± SEM of n=6 samples. ****P < 0.0001, ns > 0.05 (One way ANOVA, multiple comparison test). *PDK4* and *ANGPTL4* stimulation induced by lipid treatment were significantly reduced when combined with GSK0660, approaching control level.

### *PDK4* and *ANGPTL4* expression correlates to gastric smooth muscle dedifferentiation and the acquisition of immature features in patients with obesity

We subsequently quantified the transcript levels of *PDK4* and *ANGPTL4* in gastric smooth muscle fibers dissected and isolated from human tissue samples, which included 15 patients with obesity and 4 controls, where we had previously assessed the expression of SMC markers (figure 1; supplemental figure 10). Our analysis revealed that *PDK4* mRNA is significantly upregulated in patients with obesity compared to controls (figure 5A). Although *ANGPTL4* mRNA levels showed an increase, this change did not reach statistical significance (figure 5B). Furthermore, a significant positive correlation was observed between *ANGPTL4* and *PDK4* mRNA levels in patients with obesity, suggesting a link between their expressions (figure 5C). Conversely, an inverse correlation was found between *PDK4* mRNA and SM22 protein levels (figure 5D). Considering that alterations in smooth muscle differentiation are often associated with mesenchymal immaturity, we investigated the expression of LIX1, a specific marker and regulator of stomach mesenchymal progenitors [26,27,48]. Our analysis revealed a significant upregulation of *LIX1* mRNA in patients with obesity compared to controls (figure 5E). Additionally, in patients with obesity, *LIX1* mRNA levels demonstrated showed an inverse correlation with SM22 protein expression (figure 5F). Notably, LIX1 levels showed significant positive correlations with both *PDK4* and *ANGPTL4* mRNA expressions (figure 5G,H). These findings suggest a link between the expressions of *PDK4* and *ANGPTL4* and the immaturity of gastric smooth muscle in patients with obesity (figure 6).

**Figure 5.**
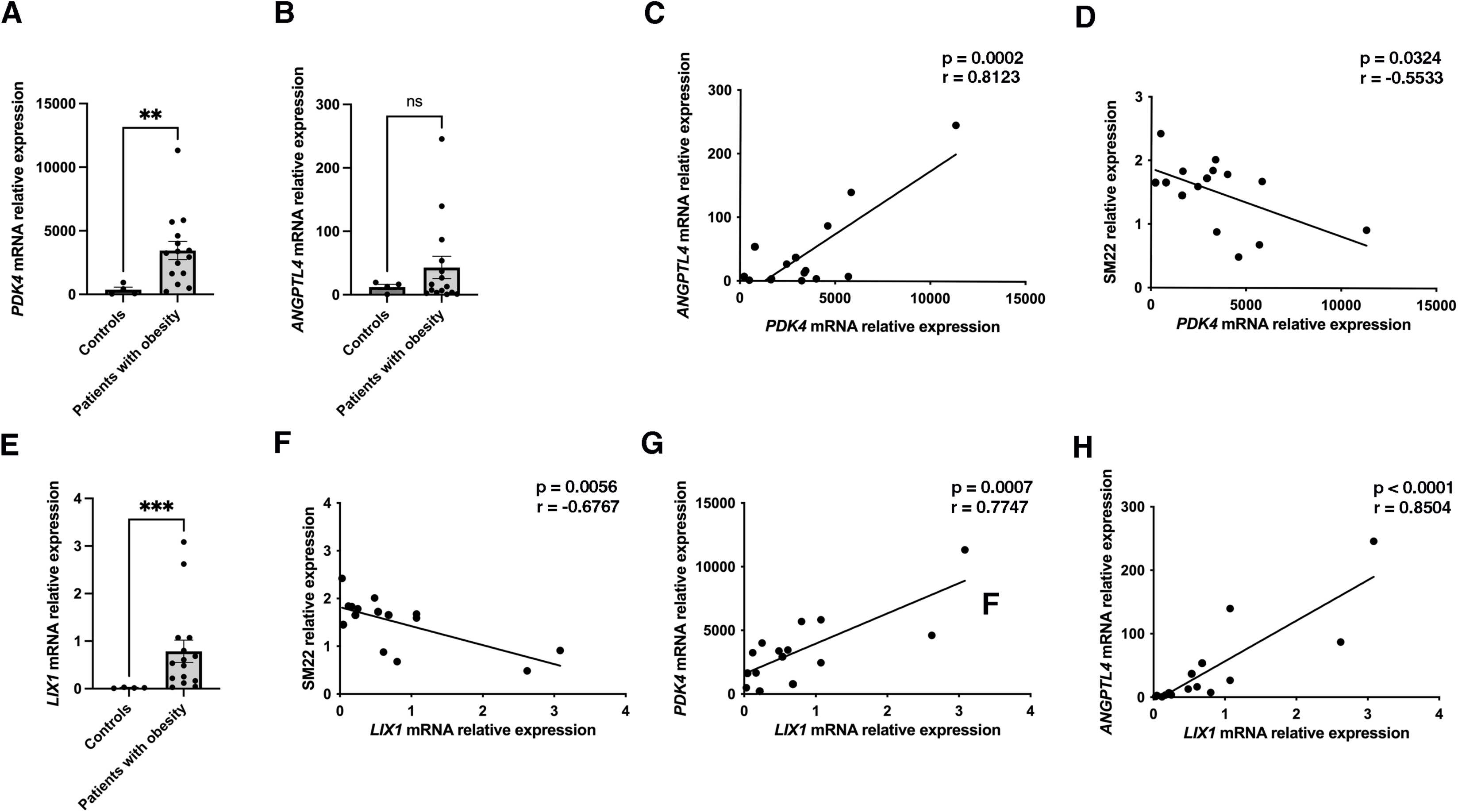
Evaluation of the expression of *PDK4*, *ANGPTL4* and *LIX1* expression and their correlation in patients with obesity. (A) Relative mRNA level of *PDK4* in gastric smooth muscle fiber extracts from adult patients with obesity (n=15) compared to controls (n=4) was assessed by RT-qPCR. Data were normalized to the levels of the housekeeping gene *HMBS*. Values are presented as mean ± SEM, and statistical analysis was performed using a two-tailed Mann– Whitney test (**P < 0.01). *PDK4* level was found to be 13-fold upregulated in patients with obesity. (B) Relative mRNA expression of *ANGPTL4* in gastric smooth muscle fiber extracts from adult patients with obesity (n=15) and controls (n=4) was evaluated by RT-qPCR. Data were normalized to *HMBS* level, presented as mean ± SEM, and analyzed using a two-tailed Mann–Whitney test (ns > 0.05). *ANGPTL4* level showed a 3.5-fold increase in patients with obesity. (C, D) The correlation between *PDK4* and *ANGPTL4* levels (C), as well as between *PDK4* level and SM22 expression (D), was analyzed using Pearson’s correlation test. P and R values are indicated on each graph. Positive correlation was observed between *PDK4* and *ANGPTL4*, while a negative correlation was identified between *PDK4* and SM22. (E) Relative mRNA level of *LIX1* in gastric smooth muscle fiber extracts from adult patients with obesity (n=15) and controls (n=4) was assessed by RT-qPCR. Data were normalized to *HMBS* level, presented as mean ± SEM, and analyzed using a two-tailed Mann–Whitney test (***P < 0.001). *LIX1* level was found to be 22-fold upregulated in patients with obesity. (F-H) The correlation between *LIX1* and SM22 expression (F), or *PDK4* (G) and *ANGPTL4* (H) levels were calculated by Pearson’s correlation test. P and R values are indicated on each graph. A negative correlation was observed between *LIX1* and SM22 expression, while positive correlations were observedbetween *LIX1* and both *PDK4* and *ANGPTL4*.

**Figure 6.**
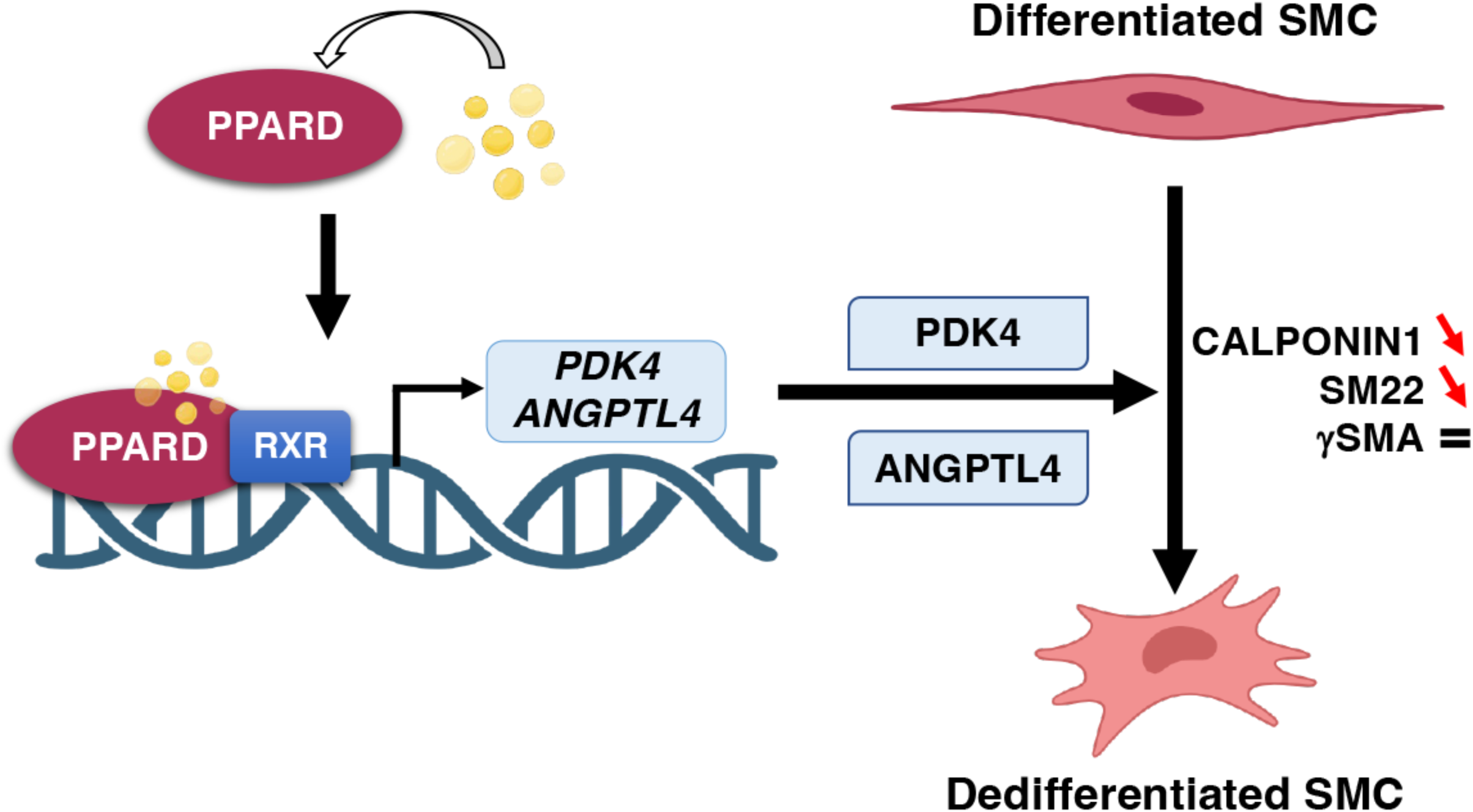
Schematic representation of the pathway leading to alterations in gastric smooth muscle in patients with obesity. Upon lipid exposure, the ligand-activated transcription factor PPARD is activated, resulting in the transcriptional upregulation of *PDK4* and *ANGPTL4*. This upregulation is crucial for initiating the dedifferentiation process of gastric SMCs observed in patients with obesity.

## DISCUSSION

The stomach plays a central role in regulating food intake through its capacity to store, mix, propel, and empty its contents, all of which require highly coordinated motor function [16]. In individuals with obesity, accelerated gastric emptying has been significantly associated with significant weight gain [7,14,15]. However, the mechanisms underlying gastric emptying dysfunction in these patients remain unclear. Gastric emptying, the process of moving stomach contents into the small intestine, is driven by peristaltic contractions—rhythmic waves of SMC activity that propel chyme toward the pylorus. Effective peristalsis requires fully differentiated SMCs for proper contractile function. We hypothesize that impaired differentiation of SMCs may contribute to the gastric emptying dysfunction observed in individuals with obesity.

In this study, we analyzed human stomach samples from patients with obesity and mouse model of high-fat diet (HFD)-induced obesity to assess the differentiation status of SMCs. Our analyses of both models revealed reduced levels of the differentiation markers SM22 and CALPONIN1 in their SMCs under obese conditions compared to controls. In contrast, the expression of γSMA, a marker of smooth muscle cell determination and identity [49], remained unchanged, indicating a specific dysregulation of the differentiation status of these cells. Additionally, we observed elevated expression of LIX1 in the gastric smooth muscle of patients with obesity, a gene typically restricted to mesenchymal progenitors during fetal development [26,27]. This finding suggests a dedifferentiation process [27,39], akin to mechanisms observed in pediatric and adult functional dysmotility syndromes, including primary visceral myopathy, inflammatory bowel disease, and achalasia [22–24,50,51]. Therefore, although gastric SMCs retain their identity, they lose their ability to contract under conditions of obesity.

The stomach is a complex organ composed of various cell types, including epithelial cells, enteric neurons, glial cells, and mesenchymal-derived cells such as SMCs and ICCs [8,16,52]. HFD-induced obesity has been shown to prevent age-associated loss in specific populations of enteric neurons, contributing to accelerated gastric emptying [36]. In the intestine, HFD-induced obesity increases the numbers and function of Lgr5+ intestinal epithelial stem cells in mammals [35]. Recent studies have investigated the immediate responses of the intestinal epithelium to HFD, revealing metabolic alterations that occur within days of exposure, leading to rapid epithelial adaptation [53,54]. Given the well-established requirement of epithelial-mesenchymal interactions in both the development and homeostasis of the GI tract [8,37,55,56], it is reasonable to hypothesize that the dedifferentiation of gastric SMCs may arise from changes in the gastric epithelium. An alternative explanation could be a direct effect of lipids on gastric SMCs. To distinguish between these two hypotheses, we first developed an *in vitro* model of differentiated human gastric SMCs that we treated with a lipid mixture enriched in oleic and palmitic acids. This mixture had previously been used to demonstrate the impact of lipids on the intestinal epithelium [35]. This treatment caused a phenotypic change in SMCs, marked by a specific reduction in differentiation markers, indicating a direct effect of lipids on SMCs. The phenotypic changes in SMCs observed *in vitro*—specifically, the loss of the differentiation markers CALPONIN1 and SM22, along with the maintenance of γSMA expression—are similar to those seen in patients with obesity and in HFD-induced mice. This highlights the possibility of a direct impact of lipids on SMCs in obese patients.

Using the previously mentioned myogenic model, we investigated the signaling mechanisms leading to SMC dedifferentiation in response to lipids. Our findings emphasized the pivotal role of a signaling cascade activated by lipid stimulation, in which PPARD regulates the mRNA levels of *PDK4* and *ANGPTL4*. PPARs are transcription factors that modulate gene expression by forming heterodimers with RXRs and binding to PPREs in the promoters of target genes, and they can be activated by various ligands, especially fatty acids. Previous research has demonstrated that PPARD is expressed during the development of the vertebrate gastrointestinal mesenchyme [57]. Our data revealed that PPARD is the most highly expressed member of the PPAR family in human gastric SMCs, and its nuclear accumulation is rapidly induced upon lipid exposure. By employing specific PPARD agonists and antagonists, we established that PPARD activation is sufficient to upregulate *PDK4* and *ANGPTL4* mRNA and is necessary for their induction following lipid treatment. This regulation correlates with the nuclear translocation of the PPARD protein, suggesting direct transcriptional control at the promoter level. These findings are consistent with studies in other cell types, such as skeletal muscle and adipocytes, indicating that PPARD may regulate *PDK4* and/or *ANGPTL4* [47,60–62]. Additionally, we observed that targeted activation of PPARD resulted in a rapid decrease in SM22 and CALPONIN1 expression, mirroring the phenotypic changes seen after lipid treatment.

Downstream of PPARD, we identified the *PDK4* and *ANGPTL4* genes as key relays in the dedifferentiation process of SMCs. PDK4, a crucial regulator of the pyruvate dehydrogenase complex, significantly contributes to obesity-related insulin resistance and metabolic dysfunction [58]. Its elevated activity in obesity enhances the formation of mitochondria-associated endoplasmic reticulum membranes while suppressing insulin signaling [58]. PDK4 is primarily expressed in skeletal muscle and the heart, with notably higher levels in the skeletal muscle of insulin-resistant individuals [63]. In patients with obesity, reduced promoter methylation of *PDK4* is observed in skeletal muscle; however, this methylation returns to levels similar to those in non-obese individuals following weight loss from Roux-en-Y gastric bypass, indicating an inverse relationship with *PDK4* mRNA expression [64]. ANGPTL4, which is essential for lipid metabolism, regulates lipoprotein lipase activity in various tissues [59]. Induced by fasting, ANGPTL4 is predominantly expressed in the liver, adipose tissue, and ischemic tissues [42,65,66]. Individuals carrying the pE40K variant of ANGPTL4 have lower triglyceride levels, higher high-density lipoprotein (HDL) cholesterol [67], and a reduced risk of coronary artery disease [68]. Moreover, these individuals demonstrate lower fasting glucose levels, improved insulin sensitivity, and a decreased likelihood of developing type 2 diabetes [69]. While numerous studies have linked serum levels of ANGPTL4 to body mass index, particularly in the context of type 2 diabetes [70], the source and functional significance of ANGPTL4 expression remain poorly understood.

PDK4 and ANGPTL4 expression had not been previously evaluated in human gastric smooth muscle. Notably, we found that *PDK4* and *ANGPTL4* transcript levels were upregulated in SMCs in response to lipid exposure or when PPARD activation was sustained in SMCs using its selective agonist, GW501516. Targeting *PDK4* or *ANGPTL4* with specific siRNAs reversed the phenotypic changes observed in SMCs following lipid stimulation, confirming their critical roles in lipid-induced SMC dedifferentiation. Importantly, we found a significant increase in *PDK4* expression in gastric smooth muscle fibers from patients with obesity compared to controls, while *ANGPTL4* levels showed a slight but non-significant increase. Furthermore, we reported a significant positive correlation between the digestive mesenchymal progenitor marker *LIX1* and both *PDK4* and *ANGPTL4* expression. Therefore, these findings may have a clinical relevance as the expression of *PDK4* and *ANGPTL4* correlates with gastric smooth muscle dedifferentiation and the development of immature features in patients with obesity. These findings emphasize the critical role of the PDK4/ANGPTL4 pathway in gastric smooth muscle cell dedifferentiation in obesity and suggest a novel mechanism that could contribute to gastric muscle dysfunction, ultimately leading to accelerated gastric emptying [6,11–14].

In conclusion, this study reveals a novel mechanistic link between obesity and gastric smooth muscle dysfunction, implicating the activation of the PPARD/PDK4/ANGPTL4 pathway. The correlation of *PDK4* and *ANGPTL4* expression with markers of mesenchymal immaturity and smooth muscle dedifferentiation underscores their potential as biomarkers and targets for therapeutic intervention in obesity-related gastric dysmotility. These insights open promising avenues for the development of targeted therapeutic strategies to mitigate the GI disorders associated with obesity.

## Acknowledgements

The authors thank the members of the “Development of visceral smooth muscle and associated pathologies” team at PhyMedExp’s institute (Montpellier) for their support and comments. We acknowledge the technical assistance provided by the personnel of MGX platform (A. Louis, Biocampus, Montpellier) and the electronic microscopy platform (C. Cazevielle, INM, Montpellier). We thank Dr. J. Ramos of her expertise in hepatogastroenterology pathology and Dr. F. Galtier for the constitution of the biological collection at Montpellier’s Hospital. We also thank Dr. L. Jaeger (PhyMedExp’s institute, Digestive Team) for her participation to the histological analysis and L. Clerat (PhyMedExp’s institute, Digestive Team) for his comment and discussion on PPAR agonist/antagonist chemistry.

## Contributors

SD, AF, KM, KK, RP, SM, FP and GW conducted the experiments, acquired and analyzed the data. LM, DN, AS, and CB provided patient samples and clinical scores. NC, PB, GW, NC, CB, SF, and PdSB designed the research studies, data compilation and analysis. PdSB and SF wrote the manuscript, with all authors participating in its editing. PdSB, and GW acquired fundings.

## Funding

This work was supported by the Agence National de la Recherche (ANR #19-CE21-0003-01, to PdSB and GW), by FIMATHO (2021-2023) and by institutional funds from INSERM, CNRS and University of Montpellier. We gratefully acknowledge the Toulouse INSERM Metatoul-Lipidomique Core Facility-MetaboHub (ANR-11-INBS-010), where lipidomic analysis were performed. SD received a PhD fellowship from the Algerian Ministry of Education. KM and SM received a PhD fellowship from the French Ministry of Education and Research (MENESR).

## Ethics approval

This study involves human participants. The biological collection was registered wih the French Ministry (numbers DC 2015-2473 and AC 2016-2760).

## Notes

### Competing Interest Statement

The authors have declared no competing interest.

## REFERENCES

1 NCD Risk Factor Collaboration (NCD-RisC). Worldwide trends in underweight and obesity from 1990 to 2022: a pooled analysis of 3663 population-representative studies with 222 million children, adolescents, and adults. Lancet. 2024;403:1027–50. doi: 10.1016/S0140-6736(23)02750-2

2 Afshin A, Reitsma MB, Murray CJL. Health Effects of Overweight and Obesity in 195 Countries. N Engl J Med. 2017;377:1496–7. doi: 10.1056/NEJMc1710026

3 De Pergola G, Silvestris F. Obesity as a major risk factor for cancer. J Obes. 2013;2013:291546. doi: 10.1155/2013/291546

4 Eslick GD. Gastrointestinal symptoms and obesity: a meta-analysis. Obes Rev. 2012;13:469–79. doi: 10.1111/j.1467-789X.2011.00969.x

5 Acosta A, Abu Dayyeh BK, Port JD, et al. Recent advances in clinical practice challenges and opportunities in the management of obesity. Gut. 2014;63:687–95. doi: 10.1136/gutjnl-2013-306235

6 Acosta A, Camilleri M. Gastrointestinal morbidity in obesity. Ann N Y Acad Sci. 2014;1311:42–56. doi: 10.1111/nyas.12385

7 Delgado-Aros S, Locke GR, Camilleri M, et al. Obesity is associated with increased risk of gastrointestinal symptoms: a population-based study. Am J Gastroenterol. 2004;99:1801–6. doi: 10.1111/j.1572-0241.2004.30887.x

8 de Santa Barbara P, van den Brink GR, Roberts DJ. Development and differentiation of the intestinal epithelium. Cell Mol Life Sci. 2003;60:1322–32. doi: 10.1007/s00018-003-2289-3

9 Steenackers N, Eksteen G, Wauters L, et al. Understanding the gastrointestinal tract in obesity: From gut motility patterns to enzyme secretion. Neurogastroenterol Motil. 2024;36:e14758. doi: 10.1111/nmo.14758

10 Chen J, Chen L, Sanseau P, et al. Significant obesity-associated gene expression changes occur in the stomach but not intestines in obese mice. Physiol Rep. 2016;4:e12793. doi: 10.14814/phy2.12793

11 Wright RA, Krinsky S, Fleeman C, et al. Gastric emptying and obesity. Gastroenterology. 1983;84:747–51.

12 Christian PE, Datz FL, Moore JG. Gastric emptying studies in the morbidly obese before and after gastroplasty. J Nucl Med. 1986;27:1686–90.

13 Acosta A, Camilleri M, Burton D, et al. Exenatide in obesity with accelerated gastric emptying: a randomized, pharmacodynamics study. Physiol Rep. 2015;3:e12610. doi: 10.14814/phy2.12610

14 Camilleri M, Malhi H, Acosta A. Gastrointestinal Complications of Obesity. Gastroenterology. 2017;152:1656–70. doi: 10.1053/j.gastro.2016.12.052

15 Pajot G, Camilleri M, Calderon G, et al. Association between gastrointestinal phenotypes and weight gain in younger adults: a prospective 4-year cohort study. Int J Obes (Lond). 2020;44:2472–8. doi: 10.1038/s41366-020-0593-8

16 Di Natale MR, Athavale ON, Wang X, et al. Functional and anatomical gastric regions and their relations to motility control. Neurogastroenterol Motil. 2023;35:e14560. doi: 10.1111/nmo.14560

17 Ward SM, Sanders KM. Physiology and pathophysiology of the interstitial cell of Cajal: from bench to bedside. I. Functional development and plasticity of interstitial cells of Cajal networks. Am J Physiol Gastrointest Liver Physiol. 2001;281:G602–611. doi: 10.1152/ajpgi.2001.281.3.G602

18 Ordög T, Ward SM, Sanders KM. Interstitial cells of cajal generate electrical slow waves in the murine stomach. J Physiol. 1999;518:257–69. doi: 10.1111/j.1469-7793.1999.0257r.x

19 Chevalier NR, Ammouche Y, Gomis A, et al. Shifting into high gear: how interstitial cells of Cajal change the motility pattern of the developing intestine. Am J Physiol Gastrointest Liver Physiol. 2020;319:G519–28. doi: 10.1152/ajpgi.00112.2020

20 Hayashi Y, Toyomasu Y, Saravanaperumal SA, et al. Hyperglycemia Increases Interstitial Cells of Cajal via MAPK1 and MAPK3 Signaling to ETV1 and KIT, Leading to Rapid Gastric Emptying. Gastroenterology. 2017;153:521–535.e20. doi: 10.1053/j.gastro.2017.04.020

21 Herring BP, Chen M, Mihaylov P, et al. Transcriptome profiling reveals significant changes in the gastric muscularis externa with obesity that partially overlap those that occur with idiopathic gastroparesis. BMC Med Genomics. 2019;12:89. doi: 10.1186/s12920-019-0550-3

22 Scirocco A, Matarrese P, Carabotti M, et al. Cellular and Molecular Mechanisms of Phenotypic Switch in Gastrointestinal Smooth Muscle. J Cell Physiol. 2016;231:295–302. doi: 10.1002/jcp.25105

23 Le Guen L, Marchal S, Faure S, et al. Mesenchymal-epithelial interactions during digestive tract development and epithelial stem cell regeneration. Cell Mol Life Sci. 2015;72:3883–96. doi: 10.1007/s00018-015-1975-2

24 Martire D, Garnier S, Sagnol S, et al. Phenotypic switch of smooth muscle cells in paediatric chronic intestinal pseudo-obstruction syndrome. J Cell Mol Med. 2021;25:4028–39. doi: 10.1111/jcmm.16367

25 Notarnicola C, Rouleau C, Le Guen L, et al. The RNA-binding protein RBPMS2 regulates development of gastrointestinal smooth muscle. Gastroenterology. 2012;143:687–697.e9. doi: 10.1053/j.gastro.2012.05.047

26 McKey J, Martire D, de Santa Barbara P, et al. LIX1 regulates YAP1 activity and controls the proliferation and differentiation of stomach mesenchymal progenitors. BMC Biol. 2016;14:34. doi: 10.1186/s12915-016-0257-2

27 Guérin A, Angebault C, Kinet S, et al. LIX1-mediated changes in mitochondrial metabolism control the fate of digestive mesenchyme-derived cells. Redox Biol. 2022;56:102431. doi: 10.1016/j.redox.2022.102431

28 Wang Z, Wang D-Z, Pipes GCT, et al. Myocardin is a master regulator of smooth muscle gene expression. Proc Natl Acad Sci USA. 2003;100:7129–34. doi: 10.1073/pnas.1232341100

29 Brittingham J, Phiel C, Trzyna WC, et al. Identification of distinct molecular phenotypes in cultured gastrointestinal smooth muscle cells. Gastroenterology. 1998;115:605–17. doi: 10.1016/s0016-5085(98)70140-4

30 Struijs M-C, Diamond IR, de Silva N, et al. Establishing norms for intestinal length in children. J Pediatr Surg. 2009;44:933–8. doi: 10.1016/j.jpedsurg.2009.01.031

31 Nair DG, Miller KG, Lourenssen SR, et al. Inflammatory cytokines promote growth of intestinal smooth muscle cells by induced expression of PDGF-Rβ. J Cell Mol Med. 2014;18:444–54. doi: 10.1111/jcmm.12193

32 Frankel D, Davies M, Bhushan B, et al. Cholesterol-rich naked mole-rat brain lipid membranes are susceptible to amyloid beta-induced damage in vitro. Aging (Albany NY). 2020;12:22266–90. doi: 10.18632/aging.202138

33 Petitfils C, Maurel S, Payros G, et al. Identification of bacterial lipopeptides as key players in IBS. Gut. 2023;72:939–50. doi: 10.1136/gutjnl-2022-328084

34 Meziat C, Boulghobra D, Strock E, et al. Exercise training restores eNOS activation in the perivascular adipose tissue of obese rats: Impact on vascular function. Nitric Oxide. 2019;86:63–7. doi: 10.1016/j.niox.2019.02.009

35 Beyaz S, Mana MD, Roper J, et al. High-fat diet enhances stemness and tumorigenicity of intestinal progenitors. Nature. 2016;531:53–8. doi: 10.1038/nature17173

36 Baudry C, Reichardt F, Marchix J, et al. Diet-induced obesity has neuroprotective effects in murine gastric enteric nervous system: involvement of leptin and glial cell line-derived neurotrophic factor. J Physiol. 2012;590:533–44. doi: 10.1113/jphysiol.2011.219717

37 Kedinger M, Duluc I, Fritsch C, et al. Intestinal epithelial-mesenchymal cell interactions. Ann N Y Acad Sci. 1998;859:1–17. doi: 10.1111/j.1749-6632.1998.tb11107.x

38 Faure S, McKey J, Sagnol S, et al. Enteric neural crest cells regulate vertebrate stomach patterning and differentiation. Development. 2015;142:331–42. doi: 10.1242/dev.118422

39 Viti F, De Giorgio R, Ceccherini I, et al. Multi-Disciplinary Insights from The First European Forum on Visceral Myopathy 2022 Meeting. Digestive Diseases and Sciences.

40 Vaes RDW, van den Berk L, Boonen B, et al. A novel human cell culture model to study visceral smooth muscle phenotypic modulation in health and disease. Am J Physiol Cell Physiol. 2018;315:C598–607. doi: 10.1152/ajpcell.00167.2017

41 Atas E, Oberhuber M, Kenner L. The Implications of PDK1-4 on Tumor Energy Metabolism, Aggressiveness and Therapy Resistance. Front Oncol. 2020;10:583217. doi: 10.3389/fonc.2020.583217

42 Sylvers-Davie KL, Davies BSJ. Regulation of lipoprotein metabolism by ANGPTL3, ANGPTL4, and ANGPTL8. Am J Physiol Endocrinol Metab. 2021;321:E493–508. doi: 10.1152/ajpendo.00195.2021

43 Konate K, Josse E, Tasic M, et al. WRAP-based nanoparticles for siRNA delivery: a SAR study and a comparison with lipid-based transfection reagents. J Nanobiotechnology. 2021;19:236. doi: 10.1186/s12951-021-00972-8

44 Neels JG, Grimaldi PA. Physiological functions of peroxisome proliferator-activated receptor β. Physiol Rev. 2014;94:795–858. doi: 10.1152/physrev.00027.2013

45 Michalik L, Wahli W. Involvement of PPAR nuclear receptors in tissue injury and wound repair. J Clin Invest. 2006;116:598–606. doi: 10.1172/JCI27958

46 Berger J, Moller DE. The mechanisms of action of PPARs. Annu Rev Med. 2002;53:409–35. doi: 10.1146/annurev.med.53.082901.104018

47 Degenhardt T, Saramäki A, Malinen M, et al. Three members of the human pyruvate dehydrogenase kinase gene family are direct targets of the peroxisome proliferator-activated receptor beta/delta. J Mol Biol. 2007;372:341–55. doi: 10.1016/j.jmb.2007.06.091

48 Guérin A, Martire D, Trenquier E, et al. LIX1 regulates YAP activity and controls gastrointestinal cancer cell plasticity. J Cell Mol Med. 2020;24:9244–54. doi: 10.1111/jcmm.15569

49 Arnoldi R, Hiltbrunner A, Dugina V, et al. Smooth muscle actin isoforms: a tug of war between contraction and compliance. Eur J Cell Biol. 2013;92:187–200. doi: 10.1016/j.ejcb.2013.06.002

50 Li C, Kuemmerle JF. Mechanisms that mediate the development of fibrosis in patients with Crohn’s disease. Inflamm Bowel Dis. 2014;20:1250–8. doi: 10.1097/MIB.0000000000000043

51 Rodrigues DM, Lourenssen SR, Kataria J, et al. Altered Esophageal Smooth Muscle Phenotype in Achalasia. J Neurogastroenterol Motil. 2024;30:166–76. doi: 10.5056/jnm23024

52 Wallace AS, Burns AJ. Development of the enteric nervous system, smooth muscle and interstitial cells of Cajal in the human gastrointestinal tract. Cell Tissue Res. 2005;319:367–82. doi: 10.1007/s00441-004-1023-2

53 Clara R, Schumacher M, Ramachandran D, et al. Metabolic Adaptation of the Small Intestine to Short- and Medium-Term High-Fat Diet Exposure. J Cell Physiol. 2017;232:167–75. doi: 10.1002/jcp.25402

54 Enriquez JR, McCauley HA, Zhang KX, et al. A dietary change to a high-fat diet initiates a rapid adaptation of the intestine. Cell Rep. 2022;41:111641. doi: 10.1016/j.celrep.2022.111641

55 Roberts DJ. Molecular mechanisms of development of the gastrointestinal tract. Dev Dyn. 2000;219:109–20. doi: 10.1002/1097-0177(2000)9999:9999<::aid-dvdy1047>3.3.co;2-y

56 McLin VA, Henning SJ, Jamrich M. The role of the visceral mesoderm in the development of the gastrointestinal tract. Gastroenterology. 2009;136:2074–91. doi: 10.1053/j.gastro.2009.03.001

57 Hojo M, Takada I, Kimura W, et al. Expression patterns of the chicken peroxisome proliferator-activated receptors (PPARs) during the development of the digestive organs. Gene Expr Patterns. 2006;6:171–9. doi: 10.1016/j.modgep.2005.06.009

58 Thoudam T, Ha C-M, Leem J, et al. PDK4 Augments ER-Mitochondria Contact to Dampen Skeletal Muscle Insulin Signaling During Obesity. Diabetes. 2019;68:571–86. doi: 10.2337/db18-0363

59 Kersten S. Role and mechanism of the action of angiopoietin-like protein ANGPTL4 in plasma lipid metabolism. J Lipid Res. 2021;62:100150. doi: 10.1016/j.jlr.2021.100150

60 Yoon JC, Chickering TW, Rosen ED, et al. Peroxisome proliferator-activated receptor gamma target gene encoding a novel angiopoietin-related protein associated with adipose differentiation. Mol Cell Biol. 2000;20:5343–9. doi: 10.1128/MCB.20.14.5343-5349.2000

61 Staiger H, Haas C, Machann J, et al. Muscle-derived angiopoietin-like protein 4 is induced by fatty acids via peroxisome proliferator-activated receptor (PPAR)-delta and is of metabolic relevance in humans. Diabetes. 2009;58:579–89. doi: 10.2337/db07-1438

62 Ordelheide A-M, Heni M, Thamer C, et al. In vitro responsiveness of human muscle cell peroxisome proliferator-activated receptor δ reflects donors’ insulin sensitivity in vivo. Eur J Clin Invest. 2011;41:1323–9. doi: 10.1111/j.1365-2362.2011.02547.x

63 McAinch AJ, Cornall LM, Watts R, et al. Increased pyruvate dehydrogenase kinase expression in cultured myotubes from obese and diabetic individuals. Eur J Nutr. 2015;54:1033–43. doi: 10.1007/s00394-014-0780-2

64 Barres R, Kirchner H, Rasmussen M, et al. Weight loss after gastric bypass surgery in human obesity remodels promoter methylation. Cell Rep. 2013;3:1020–7. doi: 10.1016/j.celrep.2013.03.018

65 Ruppert PMM, Michielsen CCJR, Hazebroek EJ, et al. Fasting induces ANGPTL4 and reduces LPL activity in human adipose tissue. Mol Metab. 2020;40:101033. doi: 10.1016/j.molmet.2020.101033

66 Kim I, Kim HG, Kim H, et al. Hepatic expression, synthesis and secretion of a novel fibrinogen/angiopoietin-related protein that prevents endothelial-cell apoptosis. Biochem J. 2000;346 Pt 3:603–10.

67 Romeo S, Pennacchio LA, Fu Y, et al. Population-based resequencing of ANGPTL4 uncovers variations that reduce triglycerides and increase HDL. Nat Genet. 2007;39:513–6. doi: 10.1038/ng1984

68 Dewey FE, Gusarova V, O’Dushlaine C, et al. Inactivating Variants in ANGPTL4 and Risk of Coronary Artery Disease. N Engl J Med. 2016;374:1123–33. doi: 10.1056/NEJMoa1510926

69 Gusarova V, O’Dushlaine C, Teslovich TM, et al. Genetic inactivation of ANGPTL4 improves glucose homeostasis and is associated with reduced risk of diabetes. Nat Commun. 2018;9:2252. doi: 10.1038/s41467-018-04611-z

70 Cinkajzlová A, Mráz M, Lacinová Z, et al. Angiopoietin-like protein 3 and 4 in obesity, type 2 diabetes mellitus, and malnutrition: the effect of weight reduction and realimentation. Nutr Diabetes. 2018;8:21. doi: 10.1038/s41387-018-0032-2

