## SupplementalLegendandTables for "Obesity induces phenotypic switching of gastric smooth muscle cells through the activation of the PPARD/PDK4/ANGPTL4 pathway"

**Supplementary information**

### Methods

#### Human tissue and mice sample preparation

Human gastric wall samples were collected and the corpus of the stomach were processed for various analyses. For histology and immunohistochemistry, samples were fixed in 4% paraformaldehyde in PBS (Corning), dehydrated with ethanol, and embedded in paraffin following standard protocols [1]. For molecular analyses, a portion of the samples underwent microdissection to isolate gastric smooth muscle fibers, which were then rapidly frozen in liquid nitrogen and stored at -80°C. These frozen samples were later used for RT-qPCR and Western Blot analyses to assess gene and protein levels. For mouse gastric wall samples, the animals were anesthetized at the end of the protocol with an intra-peritoneal injection of Dolethal (182.2 mg pentobarbital/kg body weight, Vetoquinol) and their stomachs were collected. These samples were either fixed in 10% formaldehyde for histological analysis or frozen in liquid nitrogen and stored at -80°C for biochemical assays.

#### Mouse metabolism evaluation

After 12 weeks of diet, mice from both groups underwent oral glucose tolerance tests (OGTT), and insulin tolerance tests (ITT) [2]. The OGTT was conducted at the end of the 12th week on each groups. Following an overnight fast, fasting blood glucose levels were measured using blood collected via tail clip (Caresens® N, DinnoSanteTM). The mice received a glucose solution by oral gavage (1.5g/kg), and blood glucose levels were measured at 10, 20, 30, 45, 60, 90, and 120 minutes post-administration During the same week, the other half of the group (n=7 per group) underwent an insulin tolerance test (ITT). Following a six-hour fasting period, baseline blood glucose levels were measured. Mice then received an intraperitoneal injection of insulin (1 UI/kg), and blood glucose levels were subsequently measured at 10, 20, 30, 45, 60, 90, and 120 minutes post-injection.

**Nanoparticle formulation and characterization**

The WRAP5 peptide was synthesized at the SynBio3 platform (IBMM, Montpellier) using the Fmoc strategy, following established methods [3]. Crude peptide products were purified in-house and analyzed by HPLC/MS to ensure purity greater than 95%. WRAP5 was dissolved in water (Sigma-Aldrich) to a final concentration of 400 µM (stock solution, stored at 4°C). *PDK4*- and *ANGPTL4*-siRNAs (Supplemental Table 5) and a control siRNA (si-*Neg.*, Ref.: SR-CL000-005) were purchased from Eurogentec (France) and dissolved in RNase-free water to a final concentration of 20 µM (stock solution, stored at -20°C). For the preparation of siRNA-loaded WRAP5 nanoparticle, WRAP5 and si-*Neg.* were mixed at a 20:1 molar ratio in water with 5% glucose. The nanoparticles were characterized by Dynamic Light Scattering (DLS) using a Zetasizer NanoZS (Malvern) to assess their mean size (Z-average) and distribution homogeneity (PdI). They were then stored at 4°C for 24 hours before use on human gastric smooth muscle cells (SMCs).

**Cell viability and cytotoxicity measurement**

Cell viability was assessed using with Muse® Count & Viability kit, a flow cytometry assay. Human gastric SMCs (Innoprot Innovative, Spain) [4] differentiated after 14 days of culture were treated for 3 and 7 days with lipid treatment (2% lipid mixture (Sigma; Ref. L0288) with BSA-complexed long-chain fatty acids, 30 µM palmitic acid conjugated to BSA (Bertin; Ref.

29558), and 30 µM oleic acid conjugated to BSA (Bertin; Ref. 29557)), or a 60 µM BSA control (Bertin; Ref. 29556). Cells were treated with Trypsin, resuspended in Muse® Count & Viability Reagent, and analyzed with a Muse® Cell Analyzer. Cytotoxicity of WRAP5-siRNA nanoparticles was determined using the LDH Cytotoxicity Detection KitPlus: supernatant samples were tested, with Triton X-100 serving as a positive control and a non-treated well as a negative control. Absorbance was read at 490 nm, and relative toxicity was calculated using the formula: [(experimental value – non-treated value) / (Triton value – non-treated value)] × 100.

**Transmission electron microscopy and immunofluorescence**

For transmission electron microscopy (TEM), human gastric SMCs treated with lipid treatment or BSA as control for 7 days were fixed in 2.5% glutaraldehyde in PHEM buffer (pH 7.2) at room temperature for 1 hour. Cells were then post-fixed in 0.5% osmium tetroxide in the dark for 2 hours. Following dehydration through a series of graded ethanol solutions (30% to 100%), the SMCs were embedded in EmBed 812 resin using an Automated Microwave Tissue Processor for Electron Microscopy (Leica EM AMW). Ultra-thin sections (70 nm) were cut with a Leica-Reichert Ultracut E microtome, stained with 1.5% uranyl acetate and lead citrate, and examined with a Tecnai F20 TEM at 200 kV at CoMET MRI facilities, INM, Montpellier, France. For immunofluorescence, human gastric SMCs were seeded on coverslips coated with Collagen Type I-coated (50 µg/mL per coverslips) and allowed to differentiate. The cells were treated or not with lipid mixture. They were then fixed with 4% paraformaldehyde in PBS, permeabilized with 0.1% Triton X-100, and blocked with 10% goat serum in PBS-tween 0.1%. They were then incubated with primary and secondary antibodies or stained with HCS CellMask Red 588/612 and Bodipy 493/503 for neutral lipids (Fisher Scientific, France, Ref.

11540326)[5]. A list of antibodies is provided in supplemental table 6. Nuclei were labeled with Hoechst staining. All samples were mounted in home-made Mowiol.

**RNA isolation and qPCR Analysis**

Total RNA was extracted from cell cultures with the RNeasy® Mini Kit (Qiagen) and from tissues with a Polytron homogenizer and Trizol lysis buffer (Invitrogen, Ref.: AM9738), following to the manufacturers' protocols. The extracted RNA was then reverse transcribed into complementary DNA using the Verso cDNA Synthesis Kit (Thermo Scientific). Quantitative PCRs (qPCRs) were performed using the LightCycler technology (Roche Diagnostics). PCR primers were designed based on sequences obtained from the National Library of Medicine database; detailed information can be found in supplemental table 7. Gene expression levels were quantified using the LightCycler analysis software (version 3.5) relative to standard curves. The results are presented as mean gene expression levels normalized to reference genes HMBS and YWHAZ, calculated using the $2^{-\Delta\Delta Ct}$ method.

**RNA-Seq Library preparation, sequencing and analysis**

RNA libraries were prepared from SMCs treated with or without a lipid mixture for 3 and 7 days (n=3 per condition) using the TruSeq Stranded mRNA Library Prep Kit (Illumina, ref. RS-122-2101), following the manufacturer's instructions (MGX, Biocampus, France). The libraries were validated with a Fragment Analyzer (Agilent) and quantified using the KAPA Library Quantification Kit (Roche, ref. KK4824). The libraries were then pooled in equimolar ratios and sequenced on a HiSeq2500 platform with a single-read protocol (50 nt, 1.5 lane flowcell). Image analysis and base calling were performed with Illumina HiSeq Control Software and the Real-Time Analysis component. Demultiplexing was done using Illumina's bcl2fastq 2.18

software. The quality of raw sequencing data was assessed using FastQC (Babraham Institute) and SAV (Sequencing Analysis Viewer), with potential contaminants identified via FastQ Screen (Babraham Institute). RNA-seq reads were aligned to the human genome (UCSC Hg38) using TopHat and Bowtie, incorporing gene model annotations from the UCSC database. Alignments with more than three mismatches were excluded. Read counts for each gene were generated using HTSeq-count 0.9.0.20 in union mode, and genes with fewer than 15 reads across all samples were filtered out prior to statistical analysis. Read counts were normalized with the Relative Log Expression (RLE) method in the Bioconductor package EdgeR. Differentially expressed genes were identified using EdgeR and DESeq2, with P-values adjusted for multiple testing using the Benjamini-Hochberg False Discovery Rate (FDR) method.

**Western blots**

Human gastric SMC cultures and patient-derived smooth muscle fibers were lysed using RIPA buffer, which consisted of 150 mM NaCl, 1% Triton X-100, 1% SDS, 50 mM Tris (pH 8), and a cOmplete EDTA-free protease inhibitor cocktail (Roche). Mouse stomach tissues were homogenized in a lysis buffer containing 50 mM Tris-HCl (pH 8), 2 mM EGTA, 1 mM DTT, 0.2% NP40, 0.02% SDS, and the same protease inhibitor cocktail. The lysates were incubated on ice for 1 hour, then centrifuged to collect the supernatants. Protein concentrations were measured using DC$^{TM}$ protein assay kit (Bio-Rad) according to the manufacturer's instructions. For protein analysis, 10 µg of protein from each sample were separated by 12% SDS-PAGE, transferred to nitrocellulose membranes, and subjected to Western blotting. The membranes were incubated with primary antibodies, followed by detection with infrared-labeled secondary antibodies, and imaged using the Odyssey infrared imaging system (LI-COR Biosystems). The antibodies used in this study are listed in supplemental table 4.

**Immunohistochemistry**

The paraffin-embedded tissue blocks were sectioned into 10 µm thick slices. After deparaffinization, antigen retrieval was performed using 0.01 M citrate buffer, pH 6.0, at 96°C for 30 minutes. Immunostaining was conducted following standard procedures [1]. Tissue sections were incubated overnight at 4°C with primary antibodies, which were subsequently detected with biotinylated secondary antibodies. Detection was amplified using the Streptavidin/Biotin kit (Vector) and visualized with 3,3’ diaminobenzidine (Sigma, France), followed by counterstaining with hematoxylin. Sections were rinsed and mounted using Mounting Medium (DAKO). Images were captured with a Nikon Multizoom AZ100 stereomicroscope and a Carl Zeiss AxioImager microscope. Antibodies used are listed in Supplemental Table 4.

### Supplementary tables

**Supplemental table 1.** Clinical characteristics of patients with obesity.

| | Patients without obesity (Controls) | Patients with obesity | Patients with obesity and without diabetes | Patients with obesity and with diabetes |
|---|---|---|---|---|
| Number | 4 | 15 | 6 | 9 |
| Gender, %female | 25 | 60 | 50 | 66.6 |
| Age | 56.5±9.1 | 47±11 | 42.3±15.4 | 50.6±5.9 |
| BMI | 23.4±3.3 | 42.2±5.8 | 43.4±5.8 | 41.5±6.1 |
| Blood Glucose level, $mmol^{-1}$ | nd | 6.3±1.3 | 5.3±0.4 | 7±1.3 |
| Insulinemia, mU $l^{-1}$ | nd | 15.7±9.9 | 12.4±3.6 | 17.9±12.3 |
| C peptitde, ng $ml^{-1}$ | nd | 3.5±1.5 | 3±0.6 | 3.9±1.9 |
| HbA1c % | nd | 6.4±0.9 | 5.5±0.2 | 7±0.7 |

Abbreviation : BMI, Body Mass Index; HbA1c, Hemoglobin A1c; nd, not determined. Data are presented as mean ± s.d.

### Supplementary tables

**Supplemental table 2.** Pathologies, surgery and treatment of control patients.

| | Gender | Age | BMI | Pathologies | Treatment | Surgery |
|---|---|---|---|---|---|---|
| Control#1 | M | 62 | 24.7 | Cardia carninoma | Radiotherapy | Esophagectomy |
| Control#2 | F | 46 | 26.34 | Gastroesophageal reflux and gastric fistula | n/a | Partial gastrectomy |
| Control#3 | M | 52 | 18.69 | Squamous cell carcinoma of the middle third of the esophagus | Chemotherapy and radiotherapy | Esophagectomy |
| Control#4 | M | 66 | 23.72 | Adenocarcinoma of the lower third of the esophagus | Chemotherapy (Paclitaxel) and radiotherapy | Esophagectomy |

### Supplementary tables

**Supplemental table 3.** Relative concentration (pg/mg of proteins) of fatty acid in human gastric SMCs treated with lipids for 3 and 7 days. Values (n=6) are expressed as mean±SEM; Statistical analysis was performed using Kruskal-Wallis test followed by Dunn's multiple comparisons test. *p<0.05 compared to the control.

| | **Control - 3 days** | **Lipid - 3 days** | **Control - 7 days** | **Lipid - 7 days** |
|---|---|---|---|---|
| **C16:0** | 126.5±18.27 | 157.4±29.73 | 136.6±10.99 | 175.7±14.57 |
| **C17:0** | 4.83±0.76 | 4.55±0.71 | 5.27±0.47 | 6.05±0.30 |
| **C18:0** | 133.3±20.55 | 137.3±21.70 | 153.4±12.65 | 176.5±6.707 |
| **C20:0** | 1.01±0.17 | 1.04±0.16 | 1.30±0.15 | 1.39±0.04 |
| **C23:0** | 4.09±0.71 | 5.02±0.74 | 4.84±0.48 | 9.28±0.16 * |
| **C16:1 n-7** | 12.05±2.05 | 11.63±2.45 | 11.26±1.37 | 9.032±1.05 |
| **C18:1 n-9** | 240.4±38.8 | 347.7±58.9 | 272.7±20.4 | 360.4±19.7 |
| **C18:1 n-7** | 82.16±13.35 | 74.19±11.79 | 97.93±7.51 | 86.17±3.64 |
| **C20:1 n-9** | 4.43±0.77 | 10.43±1.63 * | 8.582±0.83 | 16.62±0.50 * |
| **C22:1 n-9** | 4.17±0.81 | 5.16±0.75 | 5.97±1.09 | 7.05±0.55 |
| **C16:3** | 13.86±2.05 | 14.75±2.72 | 15.91±2.52 | 22.93±1.60 |
| **C18:2 n-6** | 5.55±0.91 | 6.80±1.24 | 5.43±0.44 | 6.48±0.39 |
| **C18:3 n-3** | 2.33±0.39 | 1.86±0.48 | 3.09±0.26 | 3.47±0.14 |
| **C20:2 n-6** | 20.06±3.35 | 21.40±3.23 | 27.72±2.42 | 30.10±0.85 |
| **C20:3 n-6** | 7.12±1.20 | 7.33±1.29 | 7.28±0.60 | 9.54±0.45 |
| **C20:4 n-6** | 48.21± 7.60 | 60.36±9.62 | 57.89±5.01 | 92.41±3.99 * |
| **C22:2 n-6** | 6.39±1.87 | 8.40±1.18 | 12.02±1.28 | 15.87±0.21 |
| **C22:5 n-3** | 5.28±0.93 | 9.09±1.59 | 4.82±0.46 | 9.31±0.39 ** |
| **C22:6 n-3** | 24.74±4.03 | 32.44±5.31 | 29.15±2.87 | 50.28±1.93 * |
| **Total FA** | 746.6±115.7 | 916.9±153.8 | 861.2±68.8 | 1089±52 |
| **Total SAFA** | 269.8±39.6 | 305.3±52.9 | 301.4±23.5 | 369.0±20.2 |
| **Total MUFA** | 343.2±55.2 | 449.1±75.4 | 396.4±30.4 | 479.3±24.1 |
| **Total PUFA** | 133.5±21.6 | 162.4±25.6 | 163.3±15.5 | 240.4±9.2 * |

Abbreviations: FA: fatty acid, SAFA: saturated fatty acid, MUFA: monounsaturated fatty acid, PUFA: polyunsaturated fatty acid.

### Supplementary tables

**Supplemental table 4.** Relative concentration (pg/mg of proteins) of ceramide and sphingomyelin in gastric human SMCs treated with lipids for 3 and 7 days. Values (n=6) are expressed as mean±SEM; Statistical analysis was performed using Kruskal-Wallis test followed by Dunn’s multiple comparisons test. *p<0.05; **p<0.01 compared to the control.

| | **Control - 3 days** | **Lipid - 3 days** | **Control - 7 days** | **Lipid - 7 days** |
|---|---|---|---|---|
| **Cer 18:1/16:0** | 984,5±108,9 | 351,3±25,6 ** | 613,2±28,5 | 182,8±10,9 ** |
| **Cer 18:1/16:1** | 14,13±1,37 | 4,82±0,37 ** | 7,08±0,31 | 2,25±0,15 ** |
| **Cer 18:1/18:0** | 193,8±19,3 | 46,50±3,50 ** | 99,70±6,03 | 19,26±0,98 ** |
| **Cer 18:1/18:1** | 10,61±1,05 | 3,34±0,24 ** | 5,99±0,30 | 2,28±0,18 ** |
| **Cer 18:1/20:0** | 52,87±5,15 | 12,74±1,03 ** | 28,59±2,32 | 5,74±0,29 ** |
| **Cer 18:1/22:0** | 353,8±34,9 | 90,60±7,46 ** | 218,4±17,3 | 44,17±2,42 ** |
| **Cer 18:1/24:0** | 1165±98 | 483,4±36,1 * | 1001±61 | 349,7±18,0 ** |
| **Cer 18:1/24:1** | 1125±109 | 460,8±33,3 ** | 867,7±52,8 | 340,6±16,0 ** |
| **Cer 18:1/26:0** | 50,70±5,55 | 13,41±1,11 * | 41,54±2,10 | 6,02±0,31 ** |
| **Cer 18:1/26:1** | 56,32±5,88 | 15,03±1,26 * | 42,81±1,97 | 6,76±0,32 ** |
| **SM 18:1/16:0** | 876,8±55,9 | 951,4±77,8 | 972,9±114,7 | 1062±147,7 |
| **SM 18:1/16:1** | 48,87±2,90 | 61,73±4,67 | 50,28±5,47 | 56,79±7,63 |
| **SM 18:1/18:0** | 88,37±6,57 | 97,33±6,65 | 96,94±9,27 | 103,4±9,9 |
| **SM 18:1/18:1** | 18,03±1,30 | 21,63±1,40 | 17,74±1,62 | 19,80±2,01 |
| **SM 18:1/20:0** | 31,77±2,18 | 34,09±2,46 | 34,85±3,79 | 37,03±3,77 |
| **SM 18:1/20:1** | 7,31±0,48 | 8,24±0,59 | 7,37±0,82 | 7,51±0,78 |
| **SM 18:1/22:0** | 127,7±8,11 | 138,2±10,5 | 144,8±17,5 | 135,8±18,4 |
| **SM 18:1/22:1** | 59,99±3,74 | 68,54±5,30 | 69,40±8,73 | 67,82±9,37 |
| **SM 18:1/24:0** | 305,4±19,6 | 289,2±20,9 | 340,6±43,1 | 300,7±40,6 |
| **SM 18:1/24:1** | 606,8±36,4 | 700,9±54,5 | 696,5±90,8 | 763,7±112,4 |
| **Total Cer** | 4006±384 | 1482±109 * | 2926±171 | 959±47 ** |
| **Total SM** | 2185±138 | 2380±184 | 2438±295 | 2560±352 |

Abbreviations : Cer: ceramide, SM: sphyngomyelin.

### Supplementary figure legends

**Supplemental figure 1** (A) Representative examples of immunohistochemistry staining with smooth muscle marker CALPONIN1 on adult controls and patients with obesity highlighting the diversity of affected gastric smooth muscle in these patients. Scale bars: 100 μm. (B) Evaluation of the CALPONIN1 expression level determined by immunohistochemistry staining of stomach sections from adult patients with obesity (n=15), and from controls (n=4). Using the IHC Profiler plugin (https://github.com/dbrant/ihc-profiler), we found that CALPONIN1 staining is statically more positive in Controls, whereas negative CALPONIN1 staining is more prominent in patients with obesity. Data are presented as the mean ± SEM (2nd way ANOVA; ns > 0.05; **P < 0.01; ****P < 0.0001).

**Supplemental figure 2** Impact of HDF containing 60% fat (230-HFD) for 12 weeks on metabolic status of male adult mice. Comparison of control water group to HFD group mice demonstrated global metabolic alteration with increase of body weight (A), adipocyte index (B), TA % (C), Fasting blood glucose (D), Total Cholesterol (E) and LDL (F). On similar way, HFD group mice present elevated oral glucose tolerance test (G) and Total caloric intake (H) compare to control water group mice. Data are presented as the mean ± SEM and Student's t test was applied (**P < 0.01; ***P < 0.001; ****P < 0.0001) (A-F).

**Supplemental figure 3** Human gastric smooth muscle cells (SMCs) were cultured on Collagen Type I plates over time to conduct to their differentiation. (A) Representative Western-blotting of human gastric SMC extracts from 1, 7, 14 and 21 days of culture probed with antibodies directed against specific smooth muscle proteins (SM22, CALPONIN1, and γSMA) and against GAPDH and VINCULIN as loading controls. (B) Quantification of Western-blot assay

(A) relative to GAPDH or VINCULIN. (C) Immunofluorescence of human gastric SMC culture after 21 days of culture stained with antibodies directed against specific smooth muscle proteins (CALPONIN1, $\alpha$SMA and FOXF1). Nuclei were visualized using Hoechst staining. Scale bars: 30 $\mu$m.

**Supplemental figure 4** (A) Venn diagram with the number of genes up-regulated and down-regulated using a significance cutoff of p < 0.01 at 3 days (upper panel) and 7 days (lower panel) post-lipid treatment compared to their respective controls. (B) Diagrams representing the individual expression of *ANGPTL4* (left panel) and *PDK4* (right panel) identified in human gastric SMC cultures with or without lipid treatment for 3 days. (C) Diagrams representing the individual expression of *ANGPTL4* (left panel) and *PDK4* (right panel) identified by RNA sequencing in human gastric SMC cultures with or without lipid treatment for 7 days.

**Supplemental figure 5** Relative mRNA expression of upregulated (A), downregulated (B) and inflammatory (C) genes identified by RNA sequencing using a significance cutoff of p < 0.01 in human gastric SMC cultures with or without lipid treatment for 3 and 7 days. Average of n=3.

**Supplemental figure 6** (A) RT-qPCR of *PDK4* (left panel), *ANGPTL4* (middle panel) and *PPARD* (right panel) relative mRNA levels in human gastric SMC cultures treated for 3 days with lipid treatment at normal concentration (Concentr. 1), and at dilution 1/4 (Concentr. 2). Data were normalized to the house-keeping *HMBS* expression. Values are the mean ± SEM of n=4 samples. ****P < 0.001; *P < 0.05; ns > 0.05 (Kruskal-Wallis test followed by Dunn's multiple comparisons test). (B) Relative mRNA expression of *PDK* members determinates by RNA sequencing from human gastric SMC cultures with or without lipid treatment for 3 and 7 days.

Average of n=3. (C) Relative mRNA expression of *ANGPTL* members determinates by RNA sequencing from human gastric SMC cultures with or without lipid treatment for 3 and 7 days. Average of n=3.

**Supplemental figure 7** Characterization of WRAP5:siRNA nanoparticles used in this study. All WRAP5:siRNA complexes were formed at room temperature using a molar ratio (R) of 20 (WRAP5:siRNA=20:1). (A) Examples of the size distribution of si-*PDK4*, si-*ANGPTL4* and si-*NEG* formulated with WRAP5 and determined by DLS technology. (B) The table indicated the mean size (Z-Ave) and particle distribution of homogeneity (Pdl) of each nanoparticles used. (C) Graphical representation of the relative cytotoxicity (%) as measured using an LDH assay after transfection with WRAP5:si-*NEG* complexes on human gastric SMCs. Non-treated cells were used as negative control (0% toxicity) whereas Triton-treated cells were used as negative control (100% toxicity). Cytotoxic condition (~20%) is not encountered in all used siRNA concentrations excepts in triton induced-cytotoxic condition (10%). (D) RT-qPCR of *PDK4* and *ANGPTL4* relative mRNA levels in human gastric SMC cultures under 3-day lipid treatment with Control siRNA (si-*NEG*) and specific siRNA directed against *PDK4* (si-*PDK4*) and *ANGPTL4* (si-*ANGPTL4*). Data were normalized to the house-keeping *HMBS* expression. Values are the mean ± SEM of n=7 samples. *P < 0.05 and **P < 0.01 (two-tailed Mann–Whitney test). *PDK4* and *ANGPTL4* stimulation are respectively diminished by 34% and 36% after 3 days of respective siRNAs under lipid treatment.

**Supplemental figure 8** (A) mRNA expression of *PPARA, PPARD and PPARG* determinates by RNA sequencing from human gastric SMC cultures with or without lipid treatment for 3 and 7 days. Average of n=3. (B) Time-course stimulation of *PDK4* and *ANGPTL4* under lipid treatment. RT-qPCR of *PDK4* (left panel), *ANGPTL4* (middle panel) and *PPARD* (right panel) relative mRNA levels in human gastric SMC cultures before treatment (0 hour) and after 6, 12 and 24 hours with lipid treatment. Values are the mean ± SEM of n=4 samples. ****P < 0.001; ***P < 0.001; **P < 0.01; *P < 0.05; ns > 0.05 (Ordinary one-way Anova test). *PDK4* stimulation is associated with linear increase, where *ANGPTL4* harbors a strong induction before a sustained statistic stimulation. *PPARD* presents only a statistical stimulation at 6 hours after lipid treatment.

**Supplemental figure 9** RT-qPCR of *PDK4* (left panel), *ANGPTL4* (middle panel) and *PPARD* (right panel) relative mRNA levels in human gastric SMC cultures treated for 3 days with or without lipid treatment, and with GW501516 (PPARD agonist, 1 μM) or DMSO as Control. Data were normalized to the house-keeping *HMBS* expression. Values are the mean ± SEM of n=5 samples. **P < 0.01; *P < 0.05; ns > 0.05 (One way ANOVA, multiple comparison test).

**Supplemental figure 10** Summary table of Two-tail Pearson's correlation test done between *PDK4, ANGPTL4, LIX1*, CALPONIN1 and SM22 expression identified in gastric smooth muscle fibers from patients with obesity classified for positive correlation (A) or for no correlation (B).

**References**


1 Rouleau C, Matécki S, Kalfa N, *et al.* Activation of MAP kinase (ERK1/2) in human neonatal colonic enteric nervous system. *Neurogastroenterol Motil*. 2009;21:207–14. doi: 10.1111/j.1365-2982.2008.01187.x

2 Meziat C, Boulghobra D, Strock E, *et al.* Exercise training restores eNOS activation in the perivascular adipose tissue of obese rats: Impact on vascular function. *Nitric Oxide*. 2019;86:63–7. doi: 10.1016/j.niox.2019.02.009

3 Konate K, Josse E, Tasic M, *et al.* WRAP-based nanoparticles for siRNA delivery: a SAR study and a comparison with lipid-based transfection reagents. *J Nanobiotechnology*. 2021;19:236. doi: 10.1186/s12951-021-00972-8

4 Guérin A, Angebault C, Kinet S, *et al.* LIX1-mediated changes in mitochondrial metabolism control the fate of digestive mesenchyme-derived cells. *Redox Biol*. 2022;56:102431. doi: 10.1016/j.redox.2022.102431

5 Qiu B, Simon MC. BODIPY 493/503 Staining of Neutral Lipid Droplets for Microscopy and Quantification by Flow Cytometry. *Bio Protoc*. 2016;6:e1912. doi: 10.21769/BioProtoc.1912
