## SupplementalFigures for "Obesity induces phenotypic switching of gastric smooth muscle cells through the activation of the PPARD/PDK4/ANGPTL4 pathway"

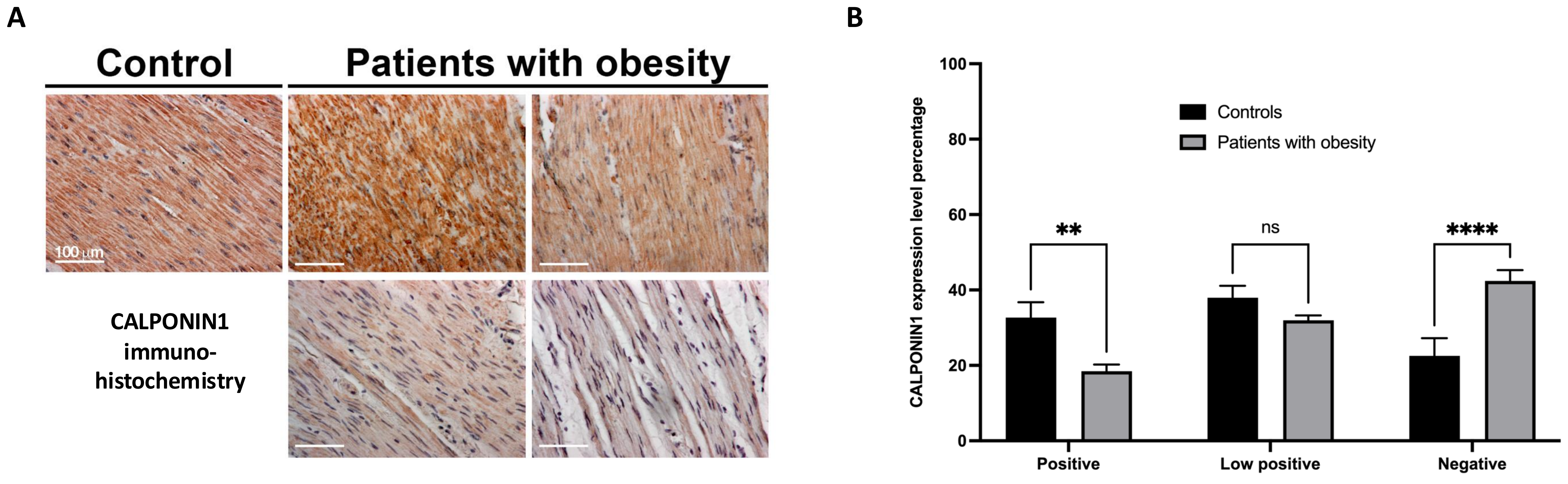


Supplemental figure 1

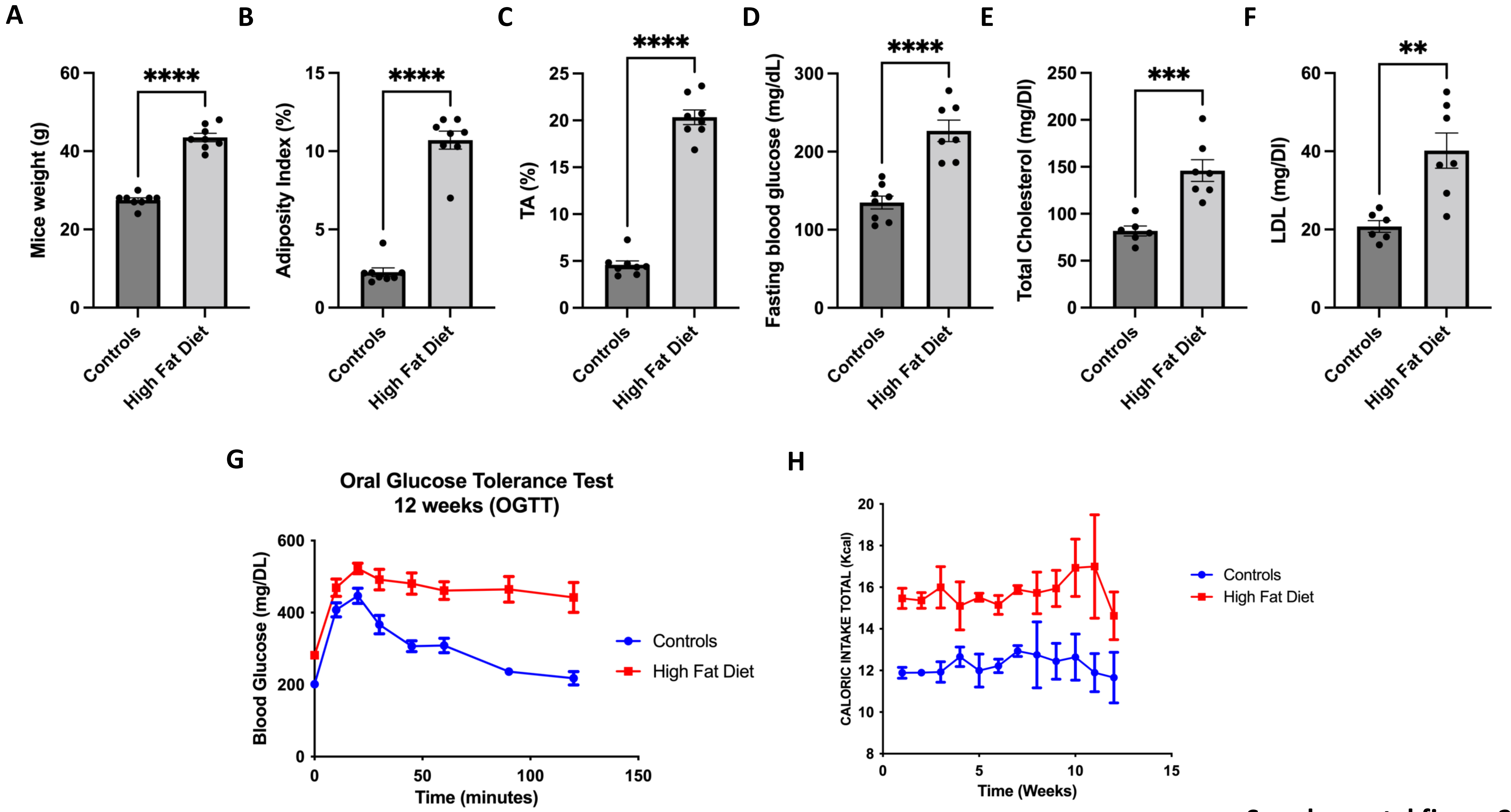


Supplemental figure 2

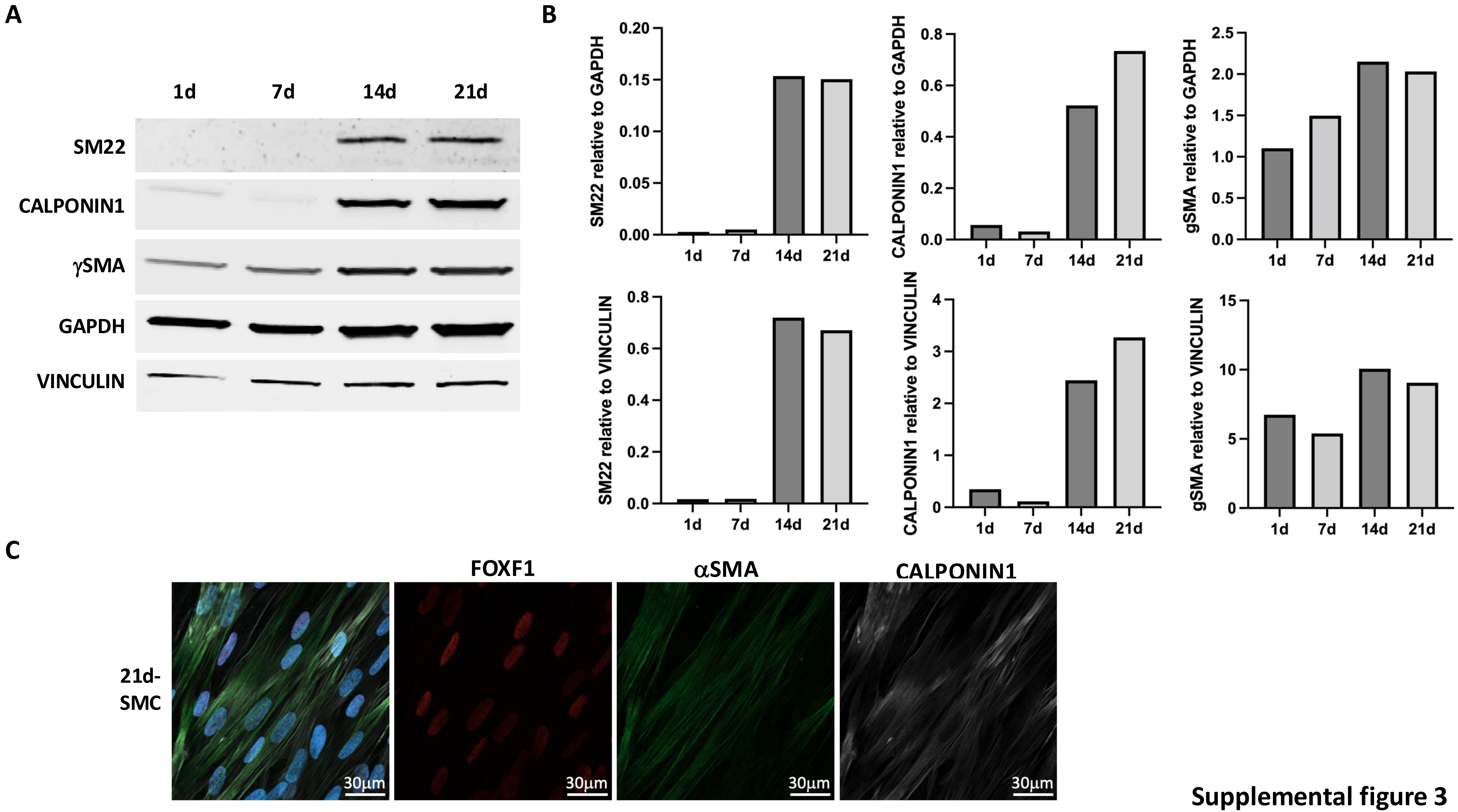


Supplemental figure 3

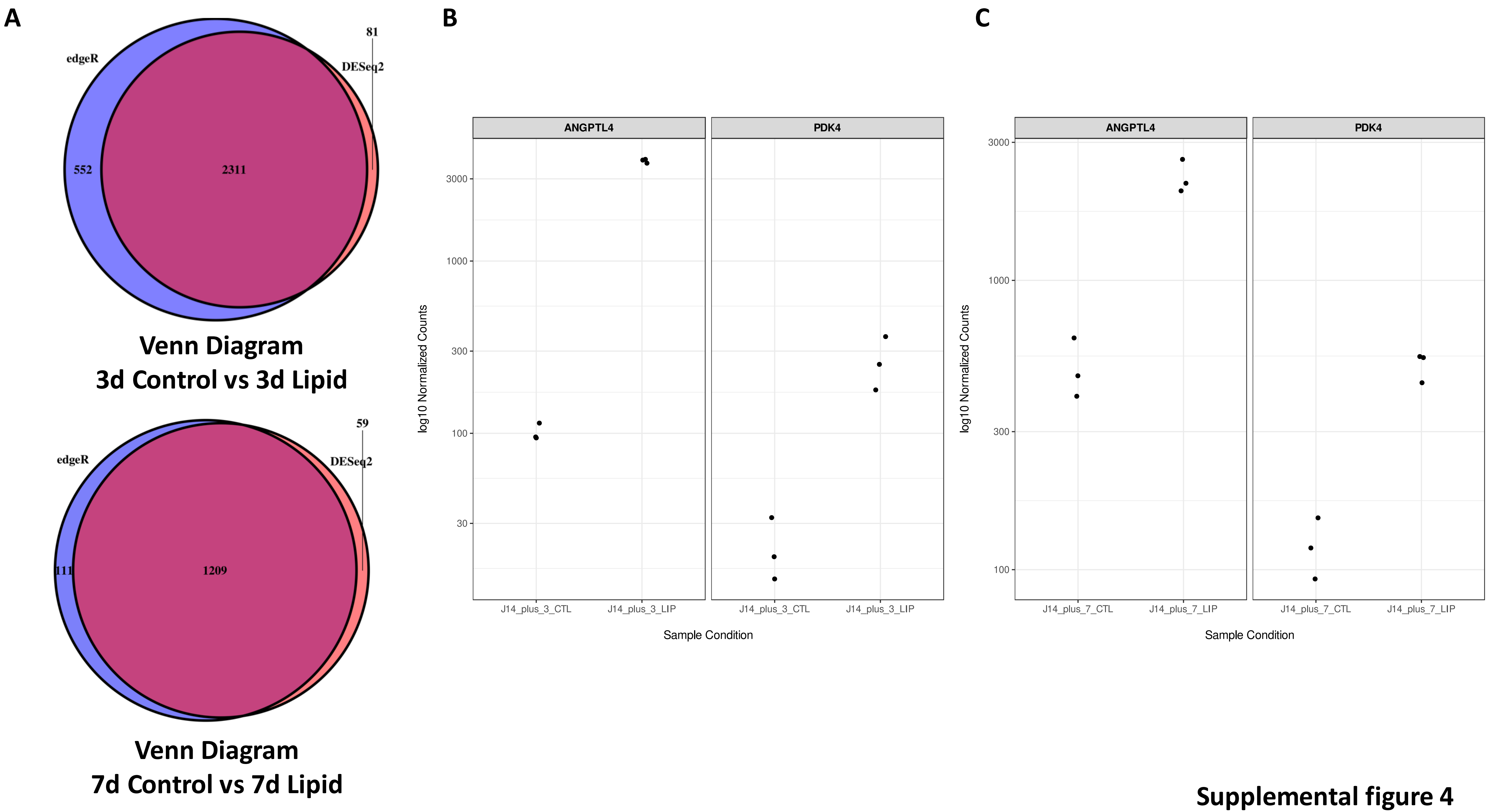


Supplemental figure 4

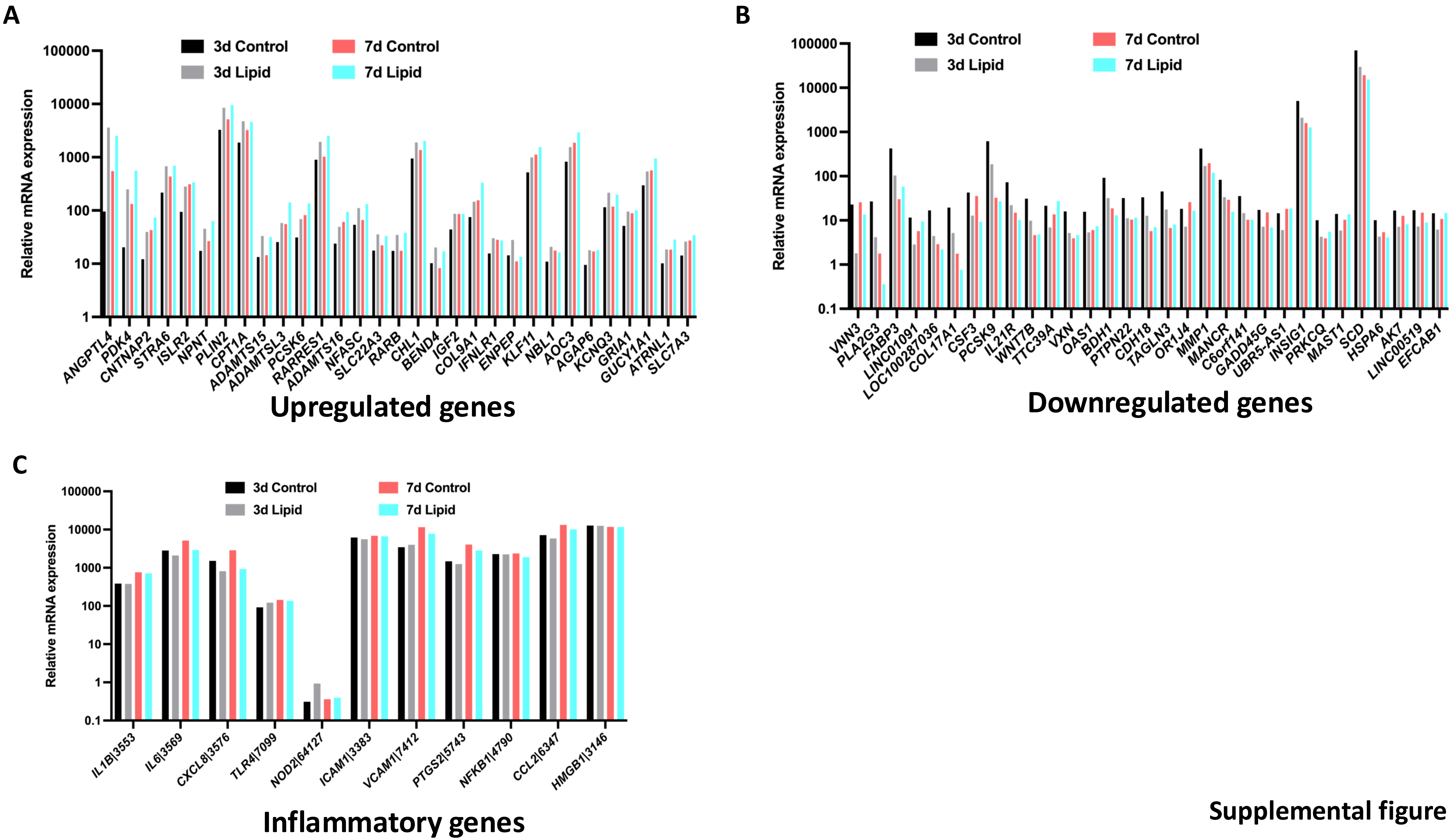


Supplemental figure 5

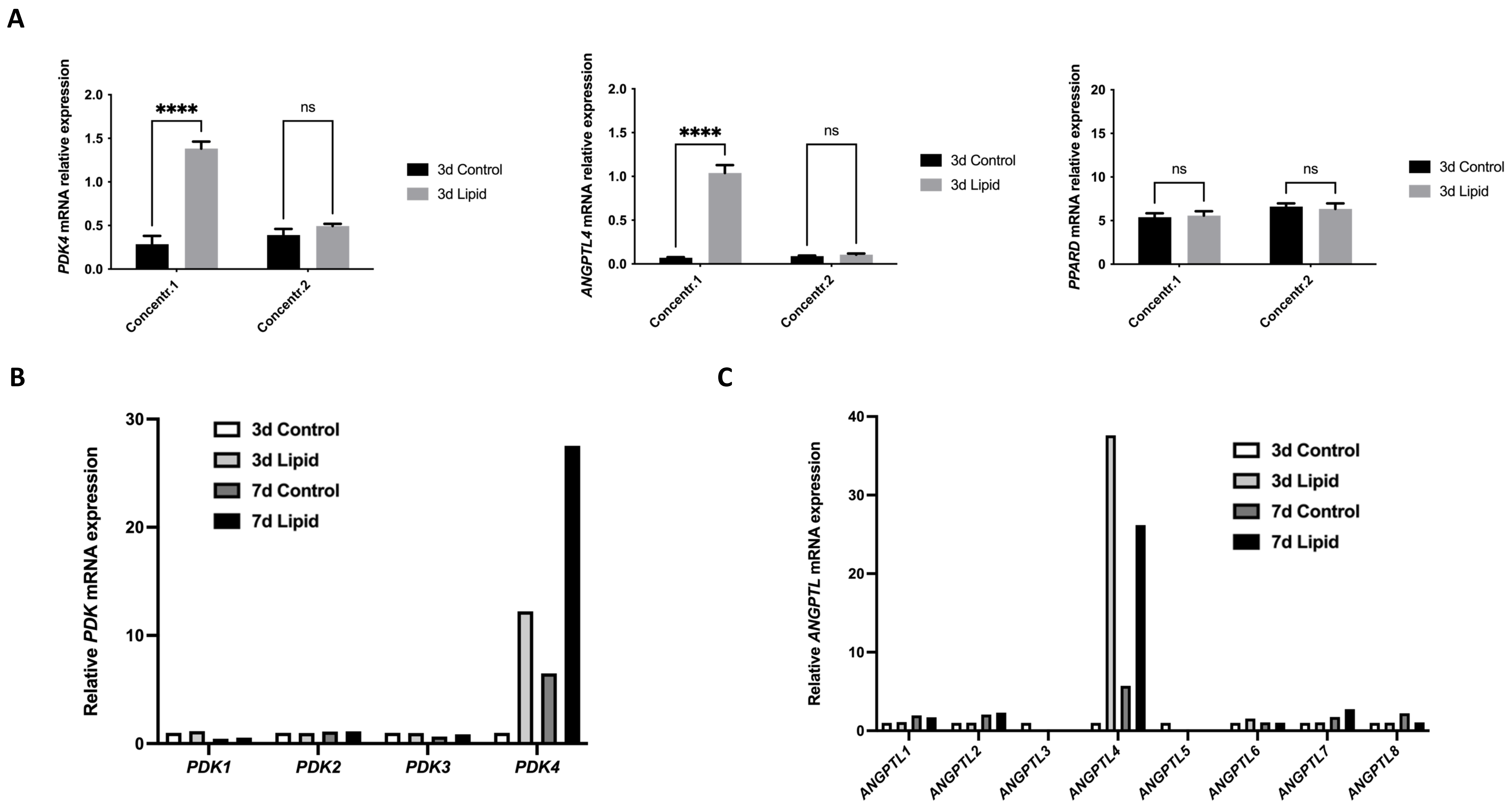


**Supplemental figure 6**

A

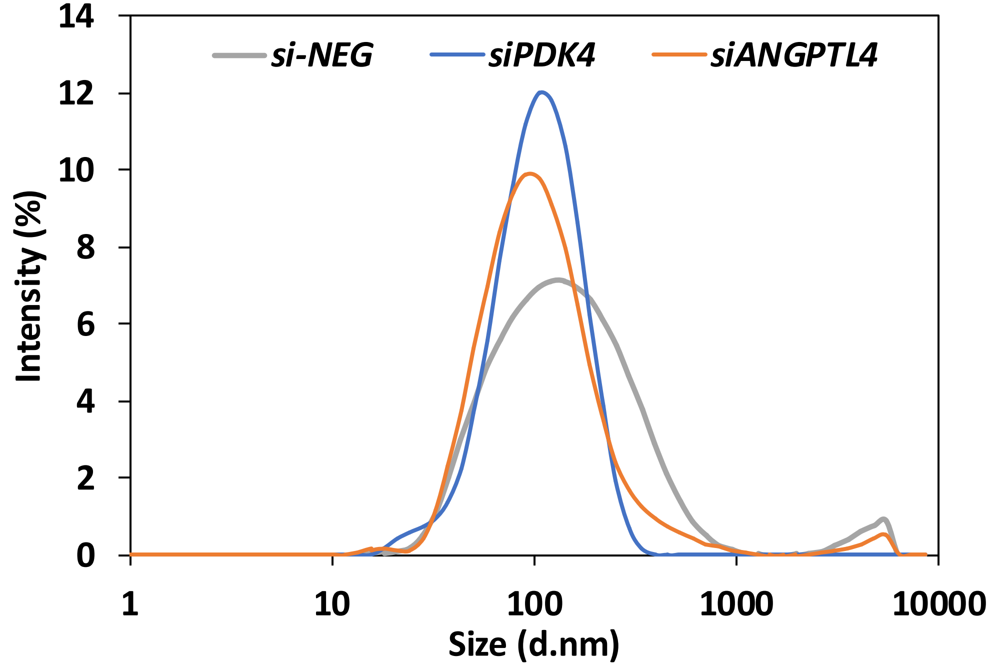


B

| Sample Name | Z-Ave (d.nm) | PdI |
|---|---|---|
| si-*NEG* | 121.9 | 0.393 |
| si-*PDK4* | 93.6 | 0.224 |
| si-*ANGPTL4* | 96.4 | 0.317 |

C

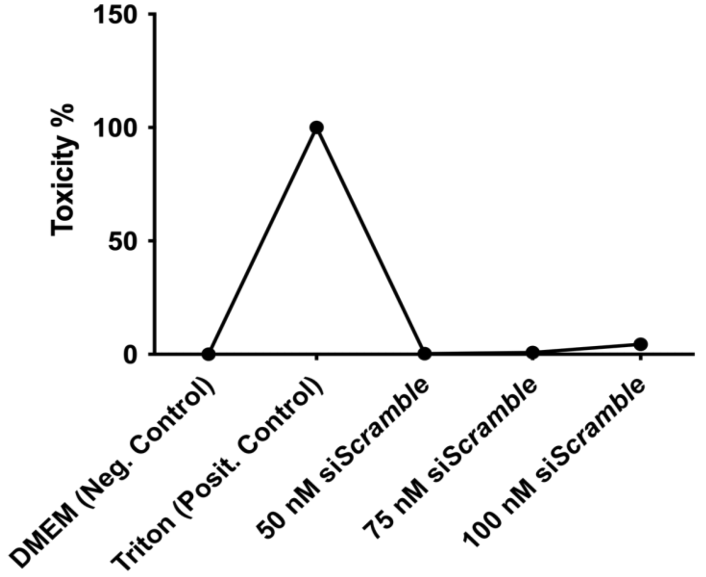


D

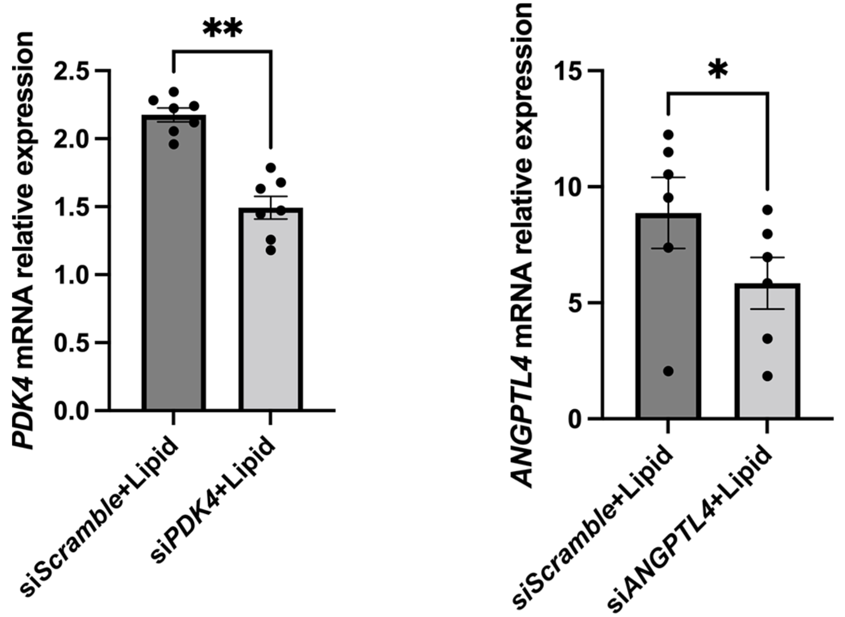


Supplemental figure 7

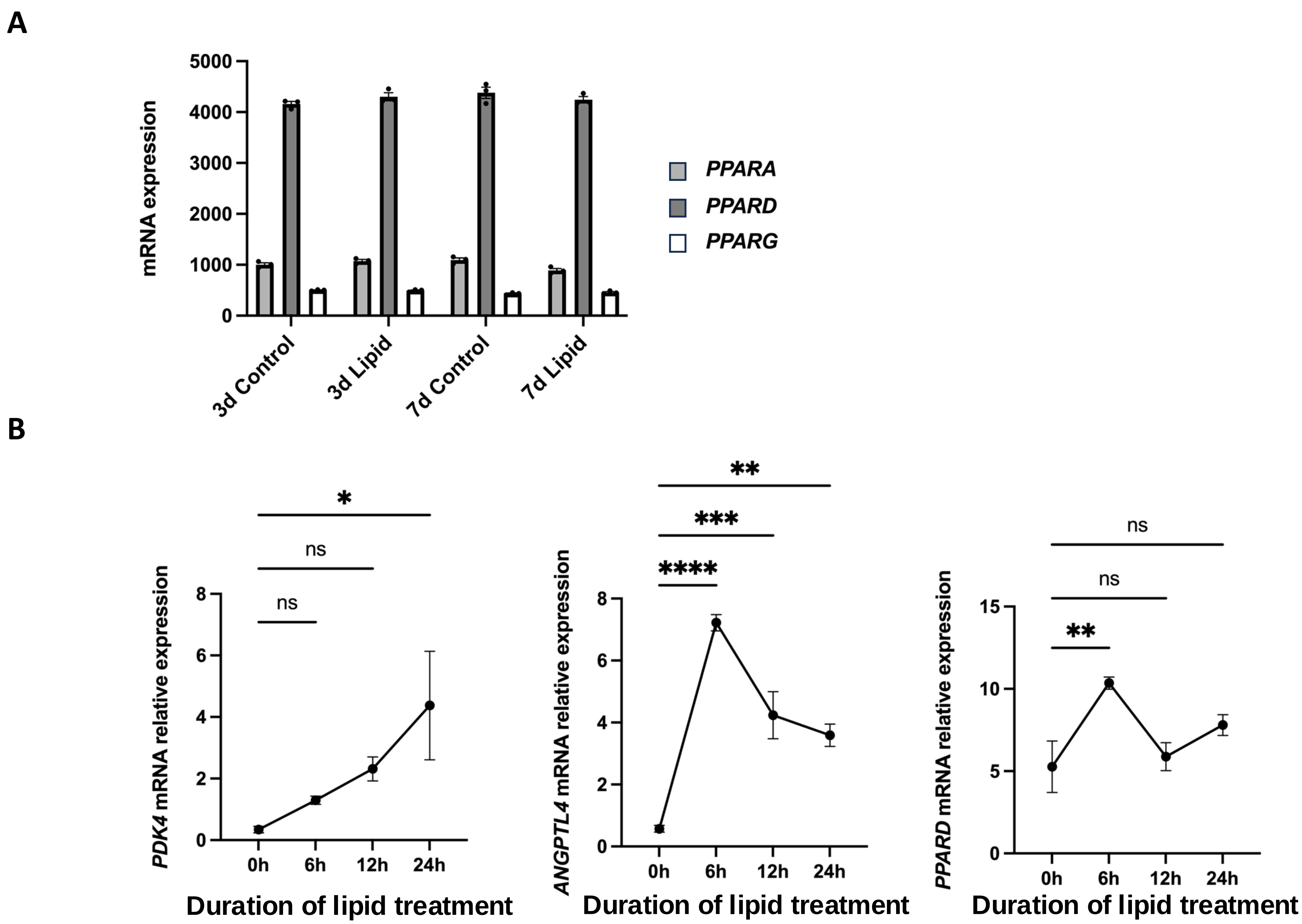


Supplemental figure 8

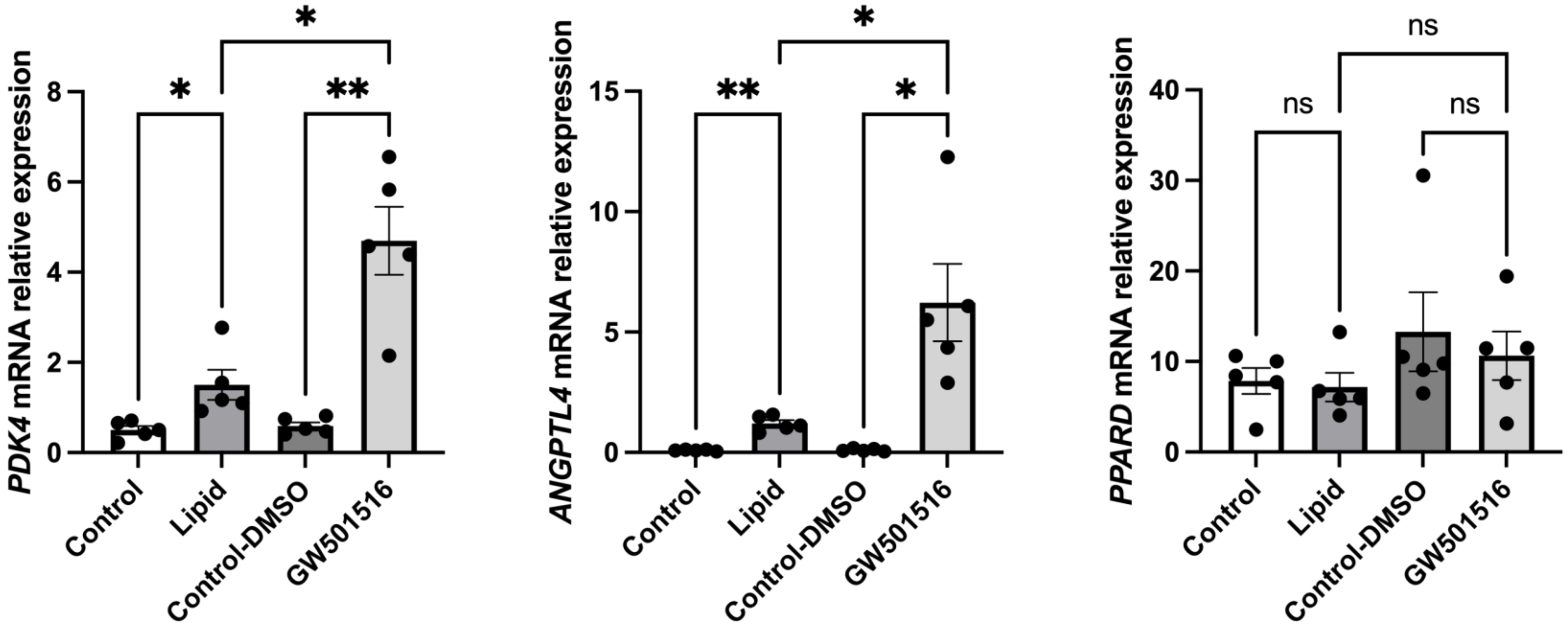


**Supplemental figure 9**

**A**

| | *PDK4* mRNA relative expression<br>vs.<br>*ANGPTL4* mRNA relative expression | *LIX1* mRNA relative expression<br>vs.<br>*ANGPTL4* mRNA relative expression | *LIX1* mRNA relative expression<br>vs.<br>*PDK4* mRNA relative expression | SM22 relative expression<br>vs.<br>*LIX1* mRNA relative expression | SM22 relative expression<br>vs.<br>*PDK4* mRNA relative expression |
|---|---|---|---|---|---|
| Pearson r | 0.8123 | 0.8504 | 0.7747 | -0.6767 | -0.5533 |
| P value | 0.0002 | <0.0001 | 0.0007 | 0.0056 | 0.0324 |
| Correlation | Elevated | Elevated | Elevated | Moderate | Moderate |

**B**

| | *PDK4* mRNA relative expression<br>vs.<br>CALPONIN relative expression | *ANGPTL4* mRNA relative expression<br>vs.<br>CALPONIN1 relative expression | *LIX1* mRNA relative expression<br>vs.<br>CALPONIN1 relative expression | *ANGPTL4* mRNA relative expression<br>vs.<br>SM22 relative expression |
|---|---|---|---|---|
| Pearson r | -0.4988 | -0.31 | -0.3115 | -0.3885 |
| P value | 0.0694 | 0.2956 | 0.2782 | 0.1524 |
| Correlation | No | No | No | No |

**Supplemental figure 10**
